# Telling mutualistic and antagonistic ecological networks apart by learning their multiscale structure

**DOI:** 10.1101/2023.04.04.535603

**Authors:** Benoît Pichon, Rémy Le Goff, Hélène Morlon, Benoît Perez-Lamarque

**Author notes:** B.P.L and H.M. jointly supervised this work.

## Abstract

1. Characterizing and understanding the processes that shape the structure of ecological networks, which represent who interacts with whom in a community, has many implications in ecology, evolutionary biology, and conservation. A highly debated question is whether and how the structure of a bipartite ecological network differs between antagonistic (e.g., herbivory) and mutualistic (e.g., pollination) interaction types.
2. Here, we tackle this question by using a multiscale characterization of network structure, machine learning tools, and a large database of empirical and simulated bipartite networks.
3. Contrary to previous studies focusing on global structural metrics, such as nestedness and modularity, which concluded that antagonistic and mutualistic networks cannot be told apart from only their structure, we find that they can be told apart by combining a meso-scale characterization of their structure and supervised machine learning. Motif frequencies appear particularly informative, with an over-representation of densely connected motifs in antagonistic networks and of motifs with asymmetrical specialization in mutualistic networks. These characteristics can be used to predict interaction types with relatively good confidence. Beyond this classical mutualism/antagonism dichotomy, we also find significant structural uniqueness linked to specific ecologies (e.g., pollination, parasitism).
4. Our results clarify structural differences between antagonistic and mutualistic networks and suggest the investigation of the structural uniqueness of specific ecologies as a promising approach for characterizing interactions beyond the coarse antagonistic/mutualistic dichotomy.

## 1. Introduction

Species in ecosystems engage in multiple types of antagonistic or mutualistic interactions, such as predation, parasitism, pollination, seed dispersal, or mycorrhizal symbioses (Bronstein, 2015; Thomas et al., 2005). Such interactions between two groups of potentially taxonomically distant species are often represented by bipartite networks (Bascompte et al., 2003; Bascompte & Jordano, 2007; Fontaine et al., 2011). Studying the structure of these interaction networks is fundamental for unraveling the different processes behind their ecological organization (Bastolla et al., 2009; Fontaine et al., 2011; Suweis et al., 2013; Thompson, 2005), understanding their evolutionary dynamics, and prioritizing conservation efforts (Bastolla et al., 2009; Olesen et al., 2007; Rezende, Lavabre, et al., 2007).

Early studies on bipartite ecological networks, focused mainly on pollination and herbivory networks, suggested that mutualistic and antagonistic networks have different structures (Bascompte et al., 2003; Thébault & Fontaine, 2008, 2010). Antagonistic networks tend to have a modular structure, where species within modules preferentially interact with each other and rarely interact with species from other modules (Krause et al., 2003; Newman, 2006b). Conversely, mutualistic networks tend to have a nested structure, where specialist species mainly interact with generalists, and generalists form a core of interactions (Jordano et al., 2002; Rohr et al., 2014; Bascompte et al., 2003). Such structural differences have been observed in both terrestrial (Fontaine et al., 2011) and marine networks (Guimarães et al., 2007). However, several studies have challenged the generality of the modular/nested dichotomy (Chagnon, 2016; Fontoura et al., 2020; Michalska-Smith & Allesina, 2019; Pilosof et al., 2014). The dichotomy seems to apply only to highly connected networks (*i.e.* with a high proportion of realized interactions), whereas networks with lower connectance are often simultaneously modular and nested (Fortuna et al., 2010). Recently, using a large and diverse dataset of empirical networks with different types of ecology (*e.g.* ant-plant, bacteria-phage, host-parasitoid,…), Michalska-Smith and Allesina (2019) showed that the structure of ecological networks is extremely heterogeneous and that global measures of network structure including nestedness and modularity are not sufficient to discriminate antagonistic and mutualistic networks (Michalska-Smith & Allesina, 2019). Song and Saavedra (2020) suggested that the confounding effect of environmental conditions indeed prevents telling antagonistic and mutualistic networks apart from such global metrics unless environmental differences are accounted for (Song & Saavedra, 2020).

Beyond the mutualistic/antagonistic dichotomy, the type of network ecology (*e.g.,* seed-dispersal compared to pollination) may significantly impact network structure. In particular, networks tend to be more modular when their interactions are more intimate, *i.e.* when the degree of biological integration between interacting individuals is high, regardless of interaction type (Fontaine et al., 2011; Guimarães et al., 2007; Hembry et al., 2018; Pires & Guimarães, 2013). For instance, within mutualistic networks, intimate ant-plant networks were found to be more modular than pollination or seed dispersal networks, the latter forming the least intimate interactions resulting in networks with low specialization (Mello et al., 2011). Within ant-plant networks, obligate symbiotic networks were found to be more specialized and modular than nectar-mediated facultative ones (Blüthgen et al., 2007; Dáttilo et al., 2013). More specific mutualistic associations between plants and mycorrhizal fungi also result in more modular mycorrhizal networks (Perez-Lamarque et al. 2022). These few examples illustrate that beyond the mutualistic/antagonistic dichotomy, the traits and ecologies of the interacting species leave an imprint on the network structures (Lewinsohn et al., 2006).

Macro-scale metrics of network structure, like nestedness and modularity, are not the only way to characterize network structure. The nestedness/modularity paradigm, as the main method to characterize bipartite network structure, has recently been questioned. Simmons, Cirtwill, et al., (2019) highlighted the limits of macro-scale metrics for capturing structural differences between bipartite networks and showed the usefulness of motifs (a small subset of species interactions exhibiting varying topologies), which capture the mesoscale structure (Milo et al., 2002). Motifs have been informative in various contexts, like the assessment of network stability, the resistance to biological invasions (Losapio et al., 2021; Milo et al., 2002; Stouffer & Bascompte, 2010; Vitali et al., 2022), or the characterization of the singular positions of functional groups (Lanuza et al., 2023). Another way to characterize the structure of interaction networks is through the spectral densities of the Laplacian graph representing each interaction network. Indeed, the eigenvectors of the Laplacian graph inform on species-level properties and can be used to identify keystone species within networks (Estrada, 2007), whereas the eigenvalues integrate information about the overall network structure, and have been linked to nestedness (Staniczenko et al., 2013) and modularity (Shen & Cheng, 2010). The spectral density of Laplacian graphs have been used in other fields, for instance, to compare structural properties of species phylogenies (Lewitus & Morlon, 2016) or brain networks (de Lange et al., 2014), but only marginally to study the structure of ecological networks (Michalska-Smith & Allesina, 2019). Since spectral densities of the Laplacian graph integrate the overall network structure, they may be informative to identify structural components that differ between antagonistic and mutualistic networks.

Here, we use a variety of network structural metrics combined with different machine learning tools to (i) investigate if there are structural components of networks, and which ones, that are distinct in mutualistic *versus* antagonistic networks, (ii) assess whether unsupervised and supervised classification approaches can robustly predict interspecific interaction type from network structure, (iii) evaluate whether currently available simulation tools for bipartite networks can help improve the accuracy of the classifiers, and (iv) assess the structural signal from specific ecologies (*e.g.,* seed-dispersal, pollination, parasitism). We compare the predictive power of the different structural metrics and classification methods. Finally, we discuss the implications of our findings for the understanding of natural communities.

## 2. Materials and methods

### Empirical interaction networks

We analyzed 343 local or regional scales bipartite interaction networks with well-known interaction types. We obtained these networks by downloading the networks from (Michalska-Smith & Allesina, 2019), among which some come from the Web of Life database (web-of-life.es (Fortuna et al., 2014)). We kept networks containing at least 10 species in each guild, as the structure of small networks is difficult to characterize (Olesen et al., 2007), in particular in terms of motif composition (preliminary results not shown). In addition, we gathered 9 mycorrhizal networks that also fit this size criterion: 6 were collected from individual papers (Jacquemyn et al., 2011, 2015; Martos et al., 2012; Montesinos-Navarro et al., 2012; Öpik et al., 2009; Xing et al., 2019) and 3 from the database generated by (Põlme et al., 2018). Among the 343 networks, 216 were mutualistic and 127 antagonistic. They comprised 9 different types of ecology: bacteria-phage (antagonistic; n=17), seed dispersal (mutualistic; n=29), host-parasite (antagonistic; n=77), host-parasitoid (antagonistic; n=10), plant-mycorrhiza (mutualistic; n=9), ant-plant (mutualistic; n=3), herbivory (antagonistic; n=23), anemone-fish (mutualistic; n=1), and pollination (mutualistic; n=174). Each network was converted into a binary matrix, taking the value 1 if species *i* interacts with species *j*, and 0 otherwise, *i.e.* we only focused on the presence/absence of interactions, not on their frequencies. The guild with the largest (resp. lowest) number of species was in rows (resp. columns).

### Comparing the structure of mutualistic *versus* antagonistic networks

Compared to the two previous studies comparing network structures between mutualistic and antagonistic networks (Michalska-Smith & Allesina, 2019; Song & Saavedra, 2020), we aimed at separately measuring the ability of different scales of network organization (macro-, meso-, and micro-scales; Fig. 1) to distinguish the structure of mutualistic *versus* antagonistic networks. We therefore considered the characterization of the structure of ecological networks by focusing on three different network scales.

**Figure 1:**
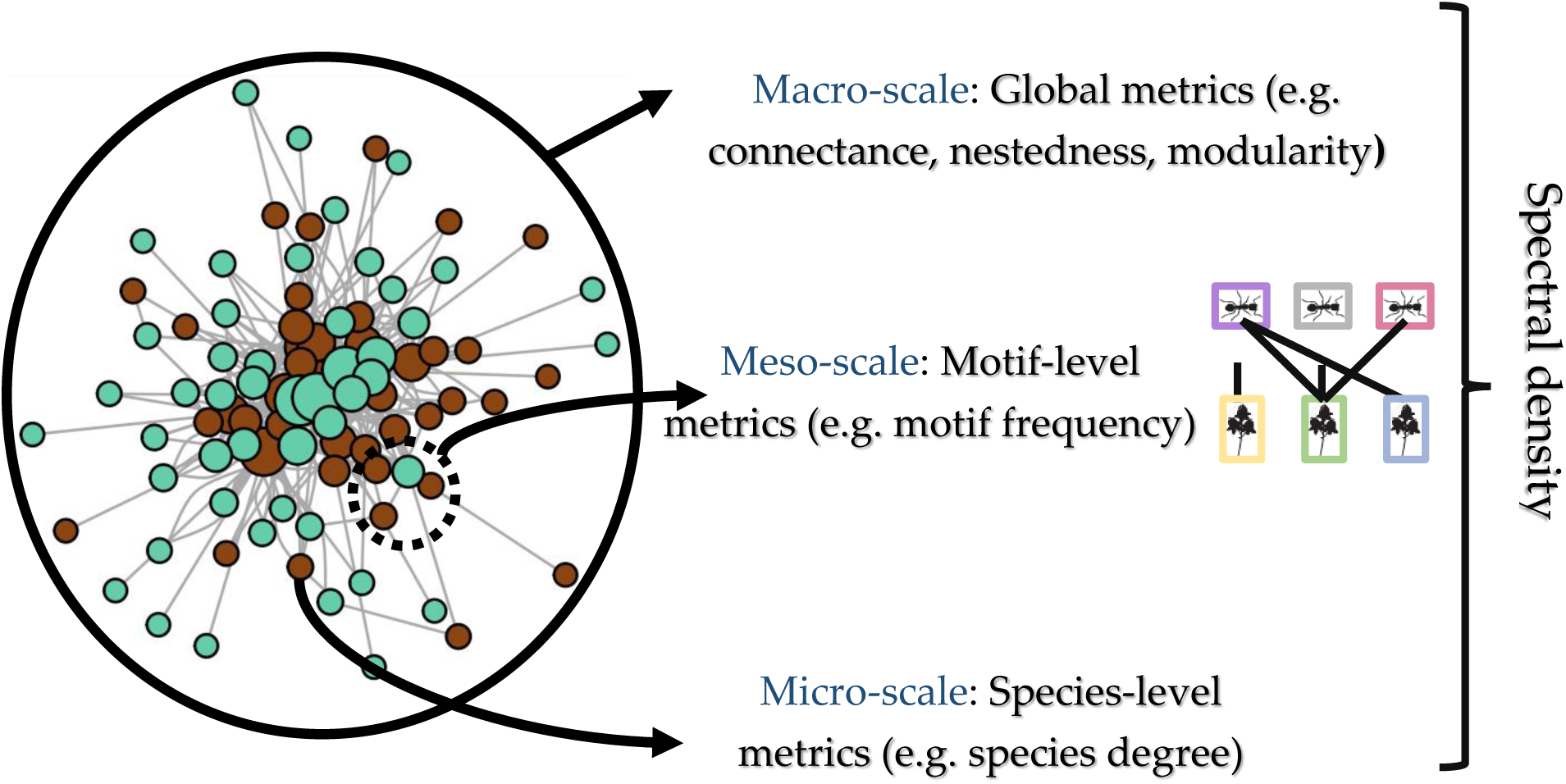
Multi-scale structure of bipartite networks. The structure of bipartite ecological networks, represented here by an ant-plant network (Blüthgen et al., 2004) can be studied at the macro-, meso-, and micro-scale. The Laplacian spectral density corresponds to a multi-scale approach from the macro to the micro-scale, whereas global metrics and motif frequencies focus on macro- and meso-scales, respectively. Blue nodes represent plant species, brown nodes ant species, and grey edges interspecific interactions.

First, we characterized the macro-scale structure of the networks using 5 global metrics: the total size (number of species in both guilds), the absolute difference in the number of species in the two guilds, the connectance, the nestedness, and the modularity. We quantified nestedness using the NODFc index that corrects for network size (“NODFc” function from the R-package maxnodf, version 1.0.0 (Hoeppke & Simmons, 2021)). We quantified modularity using the function *computeModules* from the R-package bipartite (version 2.16) (Dormann et al., 2008) with the “Beckett” method. Preliminary analyses indicated that including the total size and the absolute difference significantly improve the classification accuracies (not shown).

Second, we characterized the mesoscale structure of the networks by performing bipartite motif analyses that depict the patterns of interactions between subsets of species (called motifs; Fig. 1) (Simmons, Sweering, et al., 2019). For each network, we computed the frequencies of the 44 possible motifs involving between 2 and 6 nodes using the function *mcount* from the bmotif R-package (version 2.0.2) (Simmons et al., 2018) with the normalization procedure “normalise_sum” (*i.e.* which computes the frequency of motifs among the total number of motifs). The function *mcount* does not allow counting motifs with more than 6 nodes, and limiting the size to 6 seemed reasonable given the size of our smaller networks (10 species): the frequencies of larger motifs is likely to be more important in larger networks, independently of the type of interactions.

Third, we used a multi-scale approach stemming from graph theory that integrates the whole structural information of a network, from micro-to macro-scales: the spectral density of the Laplacian graph (Newman, 2006a). We computed the normalized Laplacian graph of each network (Newman, 2006a), defined as L_norm_ = D^-1/2^ L D^-1/2^, where D is the degree matrix (a diagonal matrix counting the number of interactions for each species) and L is the Laplacian graph, defined as the difference between the adjacency matrix (a square binary matrix whose elements represent the interactions between species of the network) and the degree matrix (Fig. S2A). The normalized Laplacian graph is a symmetric and positive-definite matrix whose eigenvalues are constrained between 0 and 2 (Banerjee & Jost, 2008), which makes them comparable between networks of different sizes, and symmetrical around 1. There are as many eigenvalues equal to 0 (or 2) as there are connected components (*i.e.* perfect modules of species that are not connected with any species from other modules), and we can thus expect a modular network to have more eigenvalues close to 0 (and 2) than a nested network. To be able to efficiently compare Laplacian graphs across networks of different sizes, we converted the eigenvalues into a spectral density using a Gaussian convolution (Lewitus & Morlon, 2016)(Fig. S2A). To do so, we adapted the *spectR* function from the RPANDA R-package (version 1.9) (Morlon et al., 2016) for bipartite networks and used a bandwidth of 0.025 for the Gaussian convolution. We tested the influence of the bandwidth value (from 0.001 to 0.091) on the resulting spectral densities. We characterized each network by extracting its spectral density values at 100 points uniformly distributed between 0 and 1.

To assess whether there are structural differences between mutualistic and antagonistic networks beyond the different types of ecology (*e.g.,* pollination, seed-dispersal, herbivory…), we controlled for their confounding effects by fitting linear mixed models (function *lmer* from the R-package lme4 (version 1.1-27)) with the type of interaction (*i.e.* mutualistic or antagonistic) as a fixed effect, the type of ecology (*e.g.,* pollination, host-parasite…) as a random effect, and one structural metric as the response variable (Type II Wald test). In addition, for each structural metric, we fitted linear models with only the type of ecology or the type of interaction as fixed effects and evaluated the best-fitting model by comparing their Akaike information criterion (AIC).

### Unsupervised classification

To visualize structural differences across antagonistic *versus* mutualistic networks, we used principal component analyses (PCA), with networks characterized either in terms of their global metrics, motif frequencies, or Laplacian spectral densities. PCAs were performed with the functions *PCA* and *fviz_pca_ind* from the FactoMineR and factoextra R-packages (versions 2.4 and 1.0.7 respectively) (Lê et al., 2008).

For each type of characterization, we then used unsupervised K-means classification on the two main principal components projections. K-means were performed with the function *kmeans* with 2 clusters, and we performed chi-squared tests to investigate whether the two clusters tend to separate mutualistic from antagonistic networks. To quantify the ability to separate antagonistic and mutualistic networks, we used a measure of classification accuracy computed as the F-score with the function *F1_score* from the R-package MLmetrics. The F-score is the harmonic mean of the precision (*e.g.,* the frequency of true mutualistic among predicted mutualistic networks) and the recall (*e.g.,* the frequency of predicted mutualistic among true mutualistic networks).

### Supervised classification

Next, we used supervised classifiers to classify antagonistic *versus* mutualistic networks based on their structure. As input for the supervised classifier, for each interaction network, we used a set of structural variables, either its global metrics, its motif frequencies, its Laplacian spectral density, or any combination of these sets of variables. As an output, the supervised classifier provides an interaction type: “mutualistic” or “antagonistic”. Supervised classifiers are based on supervised learning: the classifiers have first to be trained with a random subset of the interaction networks (the “training set”) in order to optimize its internal parameters; once trained, it can classify the other networks (the “test set”) from the input variables. We measured the accuracy of the approach using both the percentages of mutualistic and antagonistic networks from the test set that are correctly classified by the classifier and the F-scores of the classification of both types of networks.

We first investigated the accuracy of linear classifiers using logit and Lasso regressions. In Lasso regression, a constraint on the estimated model coefficients can make the regression with a high number of variables more reliable. Logit regressions were computed with the *glm* function, whereas the Lasso regressions were performed with the function *cv.glmnet* from the glmnet R-package (version 4.1-2) (Friedman et al., 2010). We found that training the classifiers with 80% of the interaction networks and testing it on the remaining 20% provided a good trade-off between the size of the test set and the accuracy of the classification.

Second, we used artificial neural networks (ANN) implemented in the *neuralnet* function from the R-package neuralnet (version 1.44.2) (Günther & Fritsch, 2010). We optimized their configuration (number of layers and number of neurons in each layer) according to the input variables. We selected a 3 layers network, constituted of the input and output layers and an intermediate layer with 5, 20, and 60 neurons for global metrics, motif frequencies, and spectral density inputs respectively. Preliminary analyses showed that these choices provided a good compromise between classification accuracy and overfitting (results not shown). We also selected a non-linear logistic transformation, and the maximal number of steps was fixed at 10^8^.

We independently trained and tested the classifiers on empirical networks using the different sets of structural input variables. For each configuration, the ANN trainings were replicated 50 times with different training and test sets, and we reported the average percentages of correct classifications among antagonistic and mutualistic networks and their respective F-scores.

### Sensitivity analyses on the supervised classification using the motif frequencies

We selected the most accurate classification approach (supervised classification using ANN on motif frequencies; see Results) and carried out a series of additional sensitivity analyses: we (i) developed a measure of classification confidence, (ii) assessed the impact of overfitting and the over-representation of mutualistic *versus* antagonistic networks and of specific types of ecology (e.g. pollination) in the training set, (iii) tested the specificity of our approach, *sensu* (Michalska-Smith & Allesina, 2019), (iv) investigated the effect of the types of ecology on our classification, and finally (v) analyzed which motif complexity is required to accurately classify networks.

First, because the training sets were limited in size (we only had a total of 343 empirical networks), we introduced a measure of the classification confidence for each empirical network. We trained and tested the ANN classifier on motif frequencies for 10,000 independent training and test sets of empirical networks: each empirical network was therefore classified on average 2,000 times with different training sets. We then computed the percentage of classifications that fell in each category, which gives an idea of classification confidence: we considered that a given network was classified as mutualistic (resp. antagonistic) with “high confidence” if it was classified as mutualistic (resp. antagonistic) in more than 75% of the classifications. Conversely, a given network was classified as mutualistic (resp. antagonistic) with “low confidence” if it was classified as mutualistic (resp. antagonistic) in more than 50% of the classifications but in less than 75% of them.

Second, we measured overfitting by comparing the percentages of correct classifications obtained on the training *versus* test sets as a function of the number of neurons in the hidden layer. In addition, we assessed the impact on our results of the over-representation of mutualistic *versus* antagonistic networks in the training set (more than 60% of the networks were mutualistic): we trained the classifier with the same number of antagonistic and mutualistic networks (from 5 to 125 networks for each type of interaction) and compared the percentages of correct classifications with the ones obtained with the same-size training sets composed of networks randomly chosen in the two interaction types. In each condition, we replicated the classification 50 times with different training sets. Similarly, we assessed the impact on our results of the over or under-representation of some types of ecology in the training set (e.g. the pollination networks represent ∼50% of the complete database). To do so, we trained the classifier with the same number of networks per type of ecology (from 5 to 20 networks for each type of ecology; if a type of ecology had fewer networks than the number per type of ecology, all networks were included in the training set) and compared the percentages of correct classifications with the ones obtained with same-size training sets composed of networks chosen randomly across ecology types.

Third, we tested the specificity of our approach *sensu* Michalska-Smith & Allesina (2019) by evaluating the classification accuracy after randomizing the network structures. To do so, we applied perturbations to the empirical networks in order to investigate whether the classification accuracies we obtained were not driven only by differences in network size, connectance, or degree distribution between antagonistic and mutualistic networks (Fig. 2A). We used two different randomization strategies, all of which maintained network size constant, and tested our classification approach on each randomized dataset. First, we randomized the networks while keeping their connectance fixed using the *shuffle.web* function of the **bipartite** R-package. Second, we randomized the networks while keeping their degree distributions fixed using the r2table algorithm implemented in the *null.model* function of the **bipartite** R-package. For each of these 2 randomization strategies, we trained the ANN using 80% of empirical networks, randomized the 20% remaining networks, and tested the ANN on these randomized networks. We replicated this randomization strategy so that each empirical network was randomized 200 times. In addition, we measured the effect of each randomization strategy on the network structure by quantifying the structural similarity between the original and randomized empirical networks. We first performed a PCA on the motif frequencies of empirical networks, and we projected randomized ones on this PCA. Then, we quantified the distance of each empirical network to the PCA-centroid of its corresponding randomized networks having either fixed connectance or degree distribution; a smaller distance indicates that the randomization strategy only slightly changes network structure. In addition, we computed the overlap between the volumes occupied by empirical networks and the randomized ones in the PCA using Jaccard similarity; a higher overlap indicates that randomized networks have kept similar motif frequencies (*hypervolume_set* in the R package **hypervolume**, v3.1.3).

**Figure 2.**
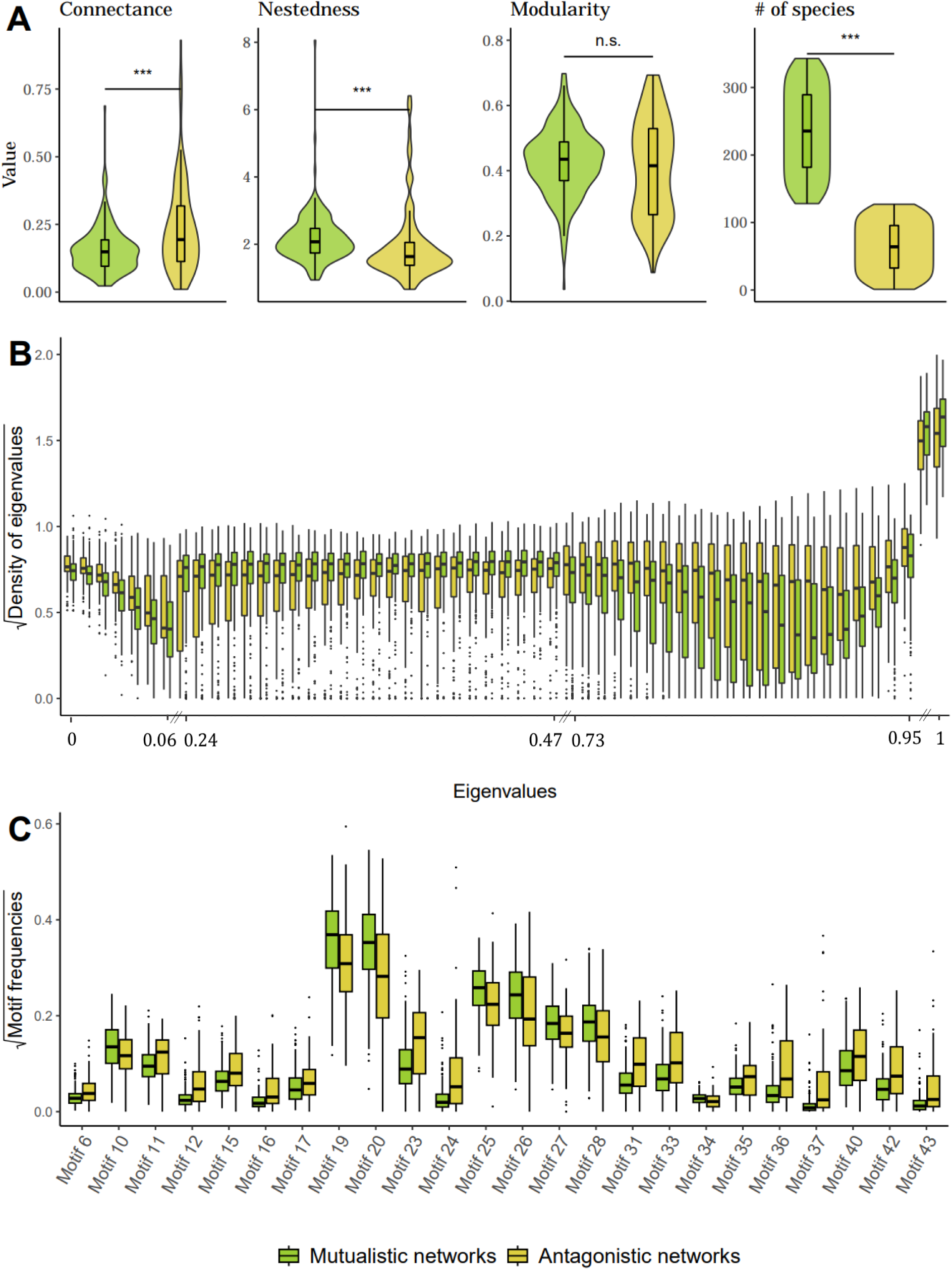
Differences between antagonistic and mutualistic networks across scales of network structure. (A) Connectance, nestedness (NODFc), and modularity (Newman’s index) of antagonistic and mutualistic empirical networks. Stars indicate the statistical significance of the difference between antagonistic and mutualistic networks, as measured by a Wilcoxon-Mann-Whitney test (*** = p-value < 0.01, ns = non-significant). (B) Densities of eigenvalues from the Laplacian spectral density that differ significantly between antagonistic and mutualistic networks (Wilcoxon-Mann-Whitney test: p-value<0.01). The discontinuities in the x-axis are due to the filtering of eigenvalues densities that differ significantly between antagonistic and mutualistic networks. (C) Frequencies of the motifs that differ significantly between antagonistic and mutualistic networks (Wilcoxon-Mann-Whitney test: p-value<0.01). Each motif is illustrated at the top of each boxplot (following the denomination of Simmons *et al*. 2018). Mutualistic networks are shown in green and antagonistic ones in yellow. Boxplots present the median surrounded by the first and third quartiles, and whiskers extend to the extreme values but no further than 1.5 of the interquartile range.

Fourth, we investigated if the classifiers were strictly learning to separate antagonistic from mutualistic networks or if they also learned the potential structural singularities of each type of ecology from our database (*i.e.,* data-linkage). We performed two different analyses to control for the signal from the type of ecology. First, we removed one by one each type of ecology from the training test (*e.g.* pollination networks), we trained the classifiers on 50 independent training sets constituted by 80% of the remaining networks, and we tested the classification on the networks from the removed type of ecology (*e.g.* are pollination networks classified among mutualistic networks when no pollination network has been used in the training?). If the classifiers learn to separate antagonistic from mutualistic networks regardless of the type of ecology, the classification for the removed type of ecology should perform as well as the original classification (*e.g.* pollination networks should be correctly classified as mutualistic networks although no pollination network has been used in the training). If they learn to classify ecologies rather than the type of interaction, then it should perform very badly (*e.g.* many pollination networks should be wrongly classified as antagonistic networks as no pollination network has been used in the training). Second, we investigated whether the structural differences between mutualistic and antagonistic networks learned by our ANN classifier were explained only by structural differences specific to each type of ecology (*e.g.,* pollination *versus* host-parasite). To do so, we shuffled the labels “mutualistic” or “antagonistic” of each type of ecology. During this shuffling process, the only constraint we imposed was keeping 5 (*resp.* 4) different types of ecology as “mutualistic” (*resp.* “antagonistic”), as in the original dataset. A shuffled dataset could for instance correspond to “antagonistic” networks represented by pollination, host-parasitoid, mycorrhizal, and anemone-fish networks, whereas “mutualistic” networks would be represented by all bacteria-phage, herbivory, host-parasite, ant-plant, and seed dispersal networks. If the structural differences from the type of ecology alone drive the ANN classifications, we expect similar accuracies with these shuffled datasets compared with the original dataset. We performed 200 of such shuffling, and for each shuffled dataset, we classified “mutualistic” *versus* “antagonistic” networks using ANN on motif frequencies with 50 different training and testing sets. In this analysis, we obtained cases where most networks were classified in one type of interaction (the most represented interaction in terms of number of networks); in this case, the percentage of correct classifications will appear very high for this interaction type, and very low for the other. Therefore, in addition to the mean accuracy of antagonistic and mutualistic networks, we measured the difference in accuracy. A large difference reflects a loss of the ANN ability to classify properly.

Finally, we investigated which level of motif complexity (size) is required to distinguish mutualistic from antagonistic networks. To do so, we measured the classification accuracy obtained when training the ANN on two subsets of all motifs, corresponding to the removal of the 6-nodes motifs, and then the removal of both 5 and 6-nodes motifs.

### Simulations of bipartite interaction networks

The accuracy of supervised classifiers often depends on the size of the available training set (i.e., data for which the classification is known), which in the case of bipartite interaction networks is rather small (n=343 here). A possible approach to increase the training set is to use simulated data. We explored this possibility with the recently-developed BipartiteEvol model (Maliet et al., 2020), an eco-evolutionary model that simulates the evolution of interacting individuals within two clades (as detailed in Supplementary Note 1).

## 3. Results

### Structure of antagonistic versus mutualistic networks

We found differences in macro-scale structures between antagonistic and mutualistic networks. For instance, nestedness was significantly higher in mutualistic networks (mean: 2.16 ± s.d. 0.71) than in antagonistic ones (1.94 ± 1.09; Wilcoxon-Mann-Whitney test: W = 8673, p-value < 0.001), whereas antagonistic networks were significantly more connected (0.23 ± 0.17) than mutualistic ones (0.15 ± 0.09; Wilcoxon-Mann-Whitney test: W = 17709, p-value < 0.001). The difference in terms of modularity was not significant (Wilcoxon-Mann-Whitney test: W = 12559, p-value = 0.19) (Fig. 2A). In addition, we found that mutualistic networks were composed of more species compared to antagonistic ones (76.1± 92.3 for antagonistic networks and 82.7 ± 72.7 for mutualistic ones; Wilcoxon-Mann-Whitney test: W = 11124, p-value = 0.003). These differences were however better explained by specific ecologies rather than the overall interaction type. Specifically, when controlling for the different types of ecology (*e.g.,* pollination, herbivory, seed dispersal…), antagonistic and mutualistic networks were not significantly different in terms of their nestedness (Wald test: χ2 = 0.060, p-value = 0.81), connectance (Wald test: χ2 = 0. 41, p-value = 0.52), modularity (Wald test: χ2 = 0.42, p-value = 0. 52), the total number of species (Wald test: χ2 = 0. 29, p-value = 0.59), and absolute difference in the number of species per guild (Wald test: χ2 = 0. 95, p-value = 0.33). Model comparisons confirmed that differences in terms of global metrics were better explained by the different types of ecology than by the antagonistic/mutualistic type of interactions (Table S1). For instance, compared to other types of ecology, bacteria-phage networks were more connected but generally less nested, while herbivory networks were more nested (Fig. S1, Table S2 A-C).

Next, using the spectral density of the Laplacian graph (Fig. S2A) on perfectly modular or nested networks revealed that modular networks have rather smooth densities with high peaks whereas nested networks have noisier densities with several small peaks (Fig. S2B-E). We found that mutualistic networks were enriched in eigenvalues near 1, while antagonistic networks had significantly more spectral eigenvalues near 0 (Fig. 2B), which was mainly driven by herbivory but not host-parasite networks (Fig. S3).

Finally, using motif frequencies, we found, following the motif denomination of (Simmons, Sweering, et al., 2019), that antagonistic networks were significantly enriched in densely connected motifs (such as motif 12 and motifs 31 to 37), whereas mutualistic networks were enriched in motifs with asymmetrical specialization, wherein specialist species tend to interact with the most generalist species in the motif (such as motifs 10, 19, 20, 25, 28 Fig. 2C). In total, 24 out of the 44 motifs differed significantly in frequency between antagonistic and mutualistic networks (p<0.01). When controlling for the type of ecology (*e.g.,* pollination *versus* herbivory), 4 motifs were significantly different in frequency between antagonistic and mutualistic networks (motifs 17, 25, 28, and 44). 31 out of 44 motifs differed significantly in frequency across the different types of ecology (Fig. S4C): for example, seed-dispersal and host-parasite networks had relatively high frequencies of 4- to 6-nodes motifs, in particular motifs 29 to 43, which may be linked to their high connectance (Figs. S1, S4., Table S2A).

### Unsupervised classifications of empirical networks

A principal component analysis (PCA) on the global metrics did not identify distinct clusters for mutualistic and antagonistic networks (Fig. 3A). Consequently, an unsupervised K-means classification on the principal component axes was not sufficient to confidently classify the networks (F-scores of 0.63 and 0.55 for mutualistic and antagonistic networks respectively) (Fig. 3A). Similarly, networks with different types of ecology (*e.g.* seed dispersal, herbivory, ant-plant…) were largely mixed on the PCA projection (Fig. S5) and did not systematically fell in one of the two clusters of the K-means classification (Table S3A).

**Figure 3.**
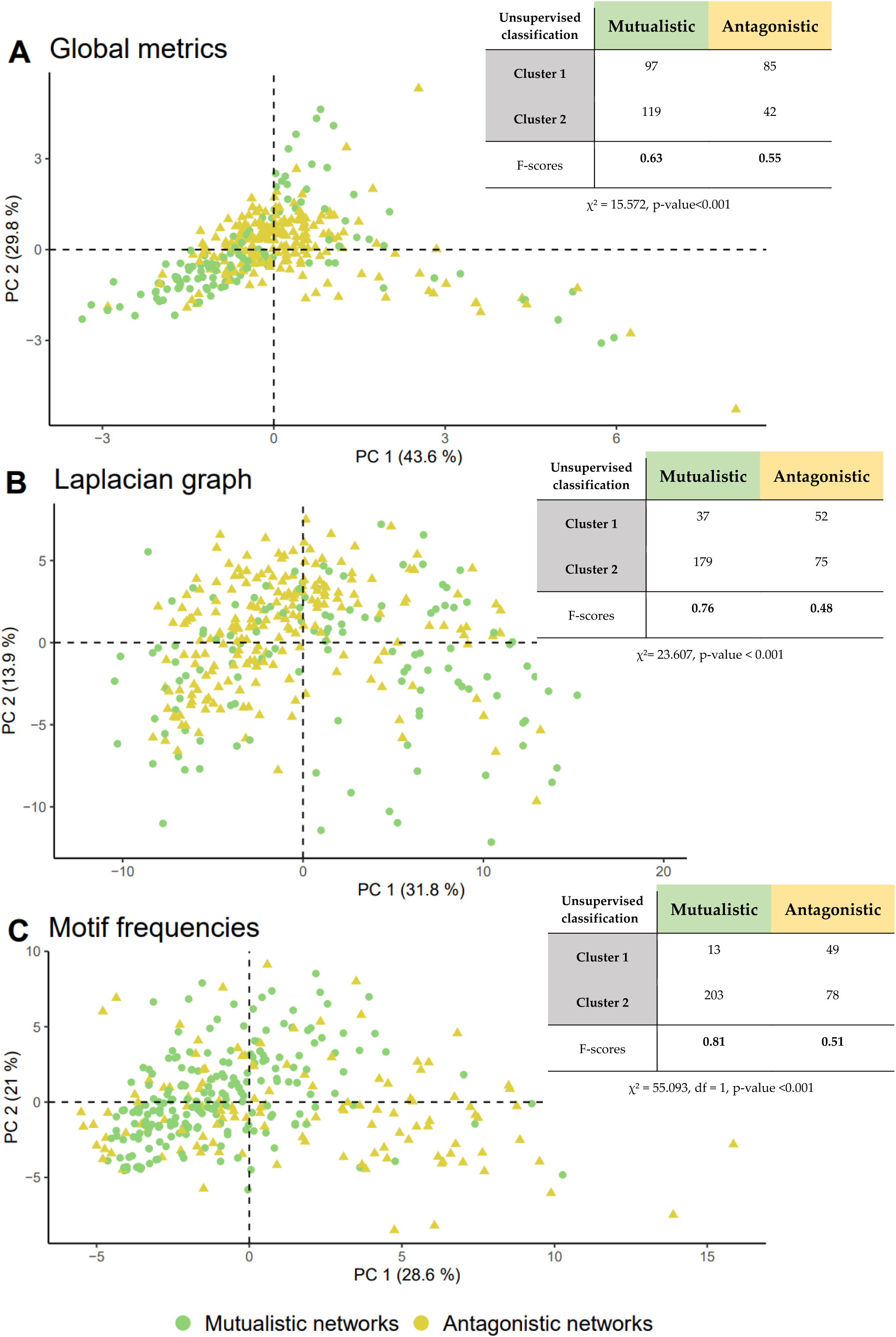
Mesoscale network structure as a scale to discriminate mutualistic *from* antagonistic networks. Projection of the 343 empirical networks on the two principal components (PC1 and PC2) obtained using principal coordinate analysis (PCA) on (A) global metrics (*i.e.* nestedness, modularity, connectance, and network size), (B) spectral density of the Laplacian graph and (C) motif frequencies. The percentage of explained variance is indicated on each axis. Each associated table corresponds to the contingency table of unsupervised K-means classification based on the global metrics (A), spectral density of the Laplacian graph (B), and motifs frequencies (C). Chi-squared test associated with the K-means classification is indicated.

PCAs performed on spectral density values (Fig. 3B) or motif frequencies (Fig. 3C) were better at separating antagonistic *versus* mutualistic empirical networks than when using the global metrics, but only slightly so (Fig. 3A). Antagonistic networks appeared to be spread on the projection, while mutualistic networks were more clustered, which is potentially linked to the high number of pollination networks in the database (Figs. S6-7). Unsupervised K-means classifications on the principal component axes were not sufficient to confidently classify mutualistic and antagonistic networks (Fig. 3B-C). Networks with different types of ecology were largely mixed between the two clusters from the K-means classification (Tables S3 B-C), suggesting that unsupervised classifications do not perform well at classifying ecological networks, regardless of the metric used to characterize their structure.

### Supervised classifications of empirical networks

We found that the two linear classifiers (Logit and Lasso regressions) mainly classified empirical networks as mutualistic, regardless of their interaction type (Table S4). Artificial neural networks (ANN) performed much better, in particular when using motif frequencies. Motif frequencies led to better classifications than the spectral density or the global metrics, and adding other variables to motif frequencies did not improve the classifications (Table S4). The ANN based on motif frequencies classified an average of 65% of the antagonistic networks (resp. 84% of the mutualistic networks) classified as antagonistic (resp. mutualistic), resulting in an F-score of 0.68 and 0.82 for antagonistic and mutualistic networks respectively (Table S4). Irrespectively of the bandwidth used to smooth the Laplacian spectrum, using the ANN with the spectral density led to lower accuracies (Tables S4-5). We therefore used ANN on motif frequencies for the remaining analyses.

### Supervised classification of the empirical networks using motif frequencies

Using our measure of classification accuracy with ANN on motif frequencies, we found that 69% (52% and 17% with high and low confidence respectively; F-score=0.74) of the antagonistic networks and 89% (76% and 13% with high and low confidence respectively; F-score=0.86) of the mutualistic networks were correctly classified (Table 1). In total, 81% (67% and 14% with high and low confidence respectively) of the networks were correctly classified (Table 1). The percentage of correct classifications varied according to the type of ecology. Indeed, most bacteria-phage, seed dispersal, and pollination networks were correctly classified by this approach (*e.g.* 23/29 of the seed dispersal networks were correctly classified, among which ∼85% with high confidence). Conversely, high intimacy mutualistic networks (*i.e.* ant-plant and mycorrhizal networks) presented frequent misclassification (only 7 out of 12 networks (∼60%) were classified as mutualistic), herbivory networks (which are low intimacy antagonistic networks) were mainly classified as mutualistic (13/23, *i.e.* 57%), and 50% (5/10) of the host-parasitoid networks were also misclassified (Table 1, Fig. S18).

**Table 1.** The type of interaction of the empirical ecological networks can be predicted based on their motif frequencies using artificial neural network classifiers.

| Network ecology | Number of networks | Prediction |  |  |  |
| --- | --- | --- | --- | --- | --- |
|  |  | Antagonistic network (high confidence) | Antagonistic network (low confidence) | Mutualistic network (high confidence) | Mutualistic network (low confidence) |
| Anemone-Fish (mutualistic) | 1 | 0 (0%) | 0 (0%) | 1 (100%) | 0 (0%) |
| Pollination (mutualistic) | 174 | 7 (4%) | 7 (4%) | 139 (80%) | 21 (12%) |
| Seed dispersal (mutualistic) | 29 | 6 (21%) | 0 (0%) | 20 (69%) | 3 (10%) |
| Ant-Plant (mutualistic) | 3 | 0 (0%) | 1 (33%) | 1 (33%) | 1 (33%) |
| Herbivory (antagonistic) | 23 | 4 (17.5%) | 6 (26%) | 9 (39%) | 4 (17.5%) |
| Mycorrhiza (mutualistic) | 9 | 3 (33.5%) | 1 (11%) | 3 (33.5%) | 2 (22%) |
| Host-Parasitoid (antagonistic) | 10 | 4 (40%) | 1 (10%) | 4 (40%) | 1 (10%) |
| Bacteria-Phage (antagonistic) | 17 | 12 (70%) | 1 (6%) | 3 (18%) | 1 (6%) |
| Host-Parasitism (antagonistic) | 77 | 46 (60%) | 14 (18%) | 10 (13%) | 7 (9%) |
| Antagonistic networks | 127 | <u>66</u> (52%) | <u>22</u> (17%) | <u>26</u> (21%) | <u>13</u> (10%) |
| Mutualistic networks | 216 | <u>16</u> (7%) | <u>9</u> (4%) | <u>164</u> (76%) | <u>27</u> (13%) |
| Correct classifications (%)<br>F-score |  | 88 (69%)<br>F-score = 0.74 |  | 191 (88%)<br>F-score = 0.86 |  |

### Robustness of the classification using motif frequencies

Overfitting had a limiting impact on the previous results (Fig. S8). In addition, controlling for the overrepresentation of mutualistic networks in the database resulted in similar mean percentage of correct classifications (Fig. S9A). Similarly, equalizing the number of networks per type of ecology in the training set did not significantly affect the overall percentages of correct classifications (Fig. S9B). Finally, we found lower classification accuracies when randomizing the empirical networks compared to the original classifications (Fig. S16a,b). Classifications were arbitrary when randomizing networks with fixed network size and connectance (mean F-scores = 0.39 and 0.53 for antagonistic and mutualistic networks respectively). When randomizing networks with fixed network size and degree distributions, classifications were not completely arbitrary (mean F-scores = 0.39 and 0.73 for antagonistic and mutualistic networks respectively (Fig. S16a,b). Such a deviation from strict random classification (50/50) can be explained as follows: mutualistic networks have a lower connectance (Fig. 2A) and are therefore more constrained during the network randomization than antagonistic networks, which results in motif frequencies closer to the original ones (Fig. S16c). Indeed, when projecting all empirical and randomized networks on a PCA, empirical mutualistic networks are significantly closer to their randomized counterparts (mean distance=8.9) compared to antagonistic networks (mean distance =14.2; Fig. S16d), and the overlap between the volumes occupied by empirical and randomized networks is higher for mutualistic networks than for antagonistic ones (Fig. S16e). In any case, the F-scores significantly dropped by 0.35 and 0.13 for antagonistic and mutualistic networks respectively when randomizing the networks, which suggests that antagonistic and mutualistic networks could not be told apart with high accuracy once randomized. Thus, differences in network size, connectance, or degree distributions between mutualistic and antagonistic networks may partially contribute to our ability to tell mutualistic and antagonistic ecological networks apart from their structure but fail to explain the high classification accuracy we report. Besides, we found that removing a given type of ecology during the ANN training decreased the percentage of correct classifications for this removed type of ecology but remained quite good in some cases (Table S7). For instance, 65% of the seed dispersal and 63% of the ant-plant networks were correctly classified when none of these networks were used during training, compared to 79% and 66% respectively when they were included (Table S7; Table 1). Also, artificially shuffling the mutualistic or antagonistic nature of each type of ecology decreased the mean percentages of correct classifications by about 10%, and the F-scores by about 0.08, while increasing the difference in percentages of correct classifications and F-scores between antagonistic and mutualistic networks by 30% and 0.3 respectively (Fig. S17). This major loss of the classification ability suggests that although the ANN classifiers partially learned the singularities of the networks linked to ecology (*i.e.* data linkage), they also learned the antagonistic/mutualistic dichotomy beyond ecology.

Finally, we found that the classification accuracy increased when including more complex motifs (Table S9). The difference was most notable between analyses carried with motifs of size at most 4 and of size at most 5 or 6 (Table S9), and rather insignificant between analyses carried with motifs of size at most 5 and at most 6. This suggests that complex motifs (with at least 5 nodes) are needed to distinguish between mutualistic and antagonistic networks, but that adding even larger motifs would not significantly increase classification accuracy.

### Ability of simulated networks to improve the accuracy of classifiers

Using simulated networks did not improved our ability to distinguish mutualistic from antagonistic networks both when using K-means classifications (results not shown) and ANN (Table S8B). Mutualistic networks, in particular, were often wrongly classified as antagonistic. When classifying simulated BipartiteEvol networks from the test sets, 70% (F-score =0.71) of antagonistic networks and 58% (F-score = 0.57) of mutualistic ones were accurately classified when using motif frequencies and ANN (Table S8A), suggesting difficulties in separating antagonistic *versus* mutualistic networks simulated with BipartiteEvol using ANN. When comparing the structure of simulated and empirical networks, we found differences in global metrics, motif composition, and density of eigen-values that may explain why simulations failed to improve the accuracy of classification (Supplementary Note 2).

## 4. Discussion

In this study, we showed that discriminating antagonistic *versus* mutualistic ecological networks based on their structure is overall possible with supervised machine learning. Artificial neural networks trained on motif frequencies reached an average of 81% correct classifications (F-scores of 0.74 and 0.86 for antagonistic and mutualistic networks respectively). Structural differences between antagonistic and mutualistic networks can be captured at the network mesoscale through differences in some motif frequencies, whereas the macro-scale modular/nested dichotomy does not seem sufficient. Beyond the classical dichotomy between mutualism/antagonism, we emphasize that the different types of ecology have an important structural imprint on the structure of ecological networks.

Using motif frequencies and ANN, we were able to correctly classify around 81% of the empirical interaction networks. This percentage of successful classifications is higher than the success rate achieved by the only previous attempt we are aware of to classify networks with ANN based exclusively on their structure (Michalska-Smith & Allesina, 2019), and achieved without accounting for environmental conditions (as in Song & Saavedra, 2020). We also demonstrated the specificity of our approach (*sensu* Michalska-Smith & Allesina, 2019): when randomizing sufficiently the network structures, we obtained lower classification accuracies, suggesting that motif frequencies, which depict patterns of species interactions at the meso-scale, capture structural differences between mutualistic and antagonistic networks that are not only due to differences in network size, connectance, or degree distributions. This therefore indicates that there are significant structural differences at the network meso-scale, which cannot be fully captured when looking at macro-scale alone using global metrics or that can be blurred when using integrated measures of network structure such as the Laplacian spectral densities.

Our analyses confirm that global metrics, including the modular/nested dichotomy, are not sufficient to discriminate between mutualistic and antagonistic network structures (Fontoura et al., 2020; Michalska-Smith & Allesina, 2019). We did not find a significant difference in modularity values between mutualistic and antagonistic networks. We did find that empirical mutualistic networks tend to be more nested and less connected than antagonistic networks, as previously suggested by (Bascompte et al., 2003; Thébault & Fontaine, 2010), but this dichotomy is not consistent enough (*i.e.,* there are too many exceptions and their variances are too large) to provide a robust classification criterion. In addition, it is worth mentioning that these global metrics are largely sensitive to the sampling effort (Nielsen & Bascompte, 2007) and may be less reliable in poorly sampled networks. In addition, there are several empirical examples of exceptions to the nested/modular dichotomy in the literature, such as a nested structure in host-parasite antagonistic networks (Pilosof et al., 2014). Within our database, some pollination and mycorrhizal networks are known to be modular (Chagnon et al., 2015; Jacquemyn et al., 2015; Martos et al., 2012; Olesen et al., 2007; Perez-Lamarque et al., 2022; Xing et al., 2019), while they assemble mainly mutualistic interactions.

Why motifs performed better than the spectral density of the Laplacian graph is unclear, as this latter representation is supposed to integrate all scales. Using a constant bandwidth value to convert the Laplacian graph spectrum into a density may result in information loss when comparing networks of different sizes. Also, transforming the discrete Laplacian spectrum into a continuous density may be problematic for small networks with only a few eigenvalues.

Motifs focus on precise structural patterns of interspecific interactions that can be missed when summarizing all the information in a single global value at the network macro-scale (Simmons, Cirtwill, et al., 2019). Although motif frequencies are highly variable across mutualistic and antagonistic networks making it impossible to accurately classify using unsupervised classification techniques (*e.g.,* PCA), there seems to exist slight differences in motif frequencies that are distinguishable using supervised machine learning. In particular, densely connected motifs (*e.g.* motifs 12 and 31 to 37) are significantly more frequent in antagonistic networks, while mutualistic networks are characterized by motifs with asymmetrical specialization (*e.g.,* motifs 10, 19, 20, 25, 28). In mutualistic networks, the emergence of motifs with asymmetrical specialization, which often parallels the emergence of nestedness, has been related to three different types of explanations. First, a pattern of asymmetric specialization may be partly explained from trait complementarity and convergence. While generalist species can establish interactions with broad range of mutualistic partners, trait complementarity generates specialization that constrains the possible interactions of specialist species (Rezende, Jordano, et al., 2007; Thompson, 2005). Second, such a pattern can emerge from differences in species abundances (neutrality hypothesis): if interactions between individuals occur at random, rare specialist species will be more likely to interact with abundant generalist species than with another rare specialist (Vázquez et al., 2009). It has been proposed that a combination of trait complementarity and abundance mechanisms best explains the observed structure of mutualistic networks (Santamaría & Rodríguez-Gironés, 2007; Vázquez et al., 2009). Finally, it has also been suggested that asymmetric specialization reduces interspecific competition (Bastolla et al., 2009; Thompson, 2005), therefore enhancing the number of coexisting species (Bascompte et al., 2006; Bastolla et al., 2009) and widening the conditions over which mutualists coexist (Rohr et al., 2014), which in the end increases the persistence and resilience of mutualistic networks to perturbations (Okuyama & Holland, 2008; Thébault & Fontaine, 2010). On the contrary, we expect antagonistic partners to engage in eco-evolutionary arm races, which may result in modules formed by highly specialized within-module interactions (i.e. densely connected motifs, which tend to be more frequent in antagonistic networks) (Maliet et al., 2020; Thompson, 2005). Such modular structures would increase the resilience of antagonistic communities (Thébault & Fontaine, 2010). Differences in motif frequencies can also be influenced by sampling biases, including missing links, preferential detection of some species, and spatial and temporal limits of the sampling. Future works will be required to identify the ecological processes and/or sampling effects that generate such structural differences between mutualistic and antagonistic networks (Brimacombe et al., 2023).

Mutualistic and antagonistic networks cover a wide range of ecologies (*e.g.* pollination, parasitism, herbivory) and our analyses showed that the type of ecology matters in the supervised classification, a phenomenon referred to as data linkage: networks with different ecologies tend to have their own structural signatures beyond the mutualistic/antagonistic dichotomy, and the ANN classifier partially learns to identify the structural uniqueness of each type of ecology. In particular, we showed that a better classification of networks of a particular type of ecology is achieved by using networks from the same ecology during the training of the classifiers. This is consistent with the result from linear mixed models that, after controlling for the type of ecology, the only significant difference between mutualistic and antagonistic networks is the frequency of a few motifs (motifs 17, 25, 28 and 44). These results on the importance of the type of ecology corroborate previous studies that found, for instance, significant structural differences between pollination and seed dispersal networks (Mello et al., 2011), between mutualistic plant-fungus and plant-animal networks (Toju et al., 2015), or across mycorrhizal networks formed by different fungal lineages (Perez-Lamarque et al., 2022). Further works are needed to better identify the species traits and/or ecological factors responsible for the structural uniqueness of the different types of network ecology. This impact of the type of ecology may also explain the low classification accuracy of some networks, like the herbivory networks, which share similar structural properties with the pollination networks in both spectral density and motifs frequency. Nevertheless, we also demonstrated by shuffling the dataset that the high classification accuracy we obtained is achieved by learning structural differences between antagonistic and mutualistic networks beyond the structural patterns specific to each type of ecology. Beyond the structural singularities of the type of ecology, there are structural differences between mutualistic and antagonistic networks that can be captured using our meso-scale approach and ANNs.

The accuracy of supervised classification approaches strongly depends on the size and quality of the dataset used for training. We used one of the largest publicly available databases of empirical networks, but it contained only 343 networks in total. In addition, it had a strong unbalanced representation of the different types of ecology, with 80% of the mutualistic networks represented by pollination networks and 60% of the antagonistic networks represented by host-parasite networks. There was also a strong imbalance in the representation of interaction intimacy, which is known to impact network structure (Fontaine et al., 2011; Guimarães et al., 2007; Hembry et al., 2018; Pires & Guimarães, 2013): mutualistic networks were mainly low intimacy networks, except ant-plants (n=3) or mycorrhizal networks (n=9), and antagonistic networks were mainly high intimacy networks, except herbivory networks (n=23). As a consequence, only ∼60% of the high-intimacy mutualistic networks and ∼40% of the low-intimacy antagonistic networks were correctly classified. These results confirm that interaction intimacy may confer a particular structure to ecological networks (Fontaine et al., 2011; Guimarães et al., 2007). Increasing the number of empirical networks, especially mutualistic networks with high intimacy (*e.g.* symbiotic networks) and antagonistic networks with low intimacy (*e.g.* trophic networks) would improve further our ability to classify ecological networks by better representing the large structural heterogeneity of ecological networks (Michalska-Smith & Allesina, 2019). In addition, using information on interaction strengths may foster our classification accuracy, as it contains relevant structural information (*e.g.,* on nestedness, Staniczenko et al., 2013). We could not test this properly as only 69 out of 343 networks in our database contained interaction strengths but the ongoing sampling of more and more weighted networks paves the way for the inclusion of interaction strengths into classification approaches in the future.

The accuracy of supervised classification approaches can also strongly depend on the type, architecture, and tuning of the classifier. For example, the overfitting we noticed in the ANN (which did not seem to negatively impact our results) could potentially be avoided by using other regularization technics, such as dropout (Srivastava et al., 2014). We chose to use simple, few-layers neural networks implemented in R for their availability and ease of usage to the large community of R users, as well as because our dataset for training was rather small. However, several other machine learning tools exist and could be tested, although they may require larger training sets; graph neural networks in particular have been specifically designed to learn information from graphs such as ecological networks (Wu et al., 2021).

One approach to better train the ANNs would be to increase the size of the training data thanks to synthetic data, obtained through either simulations or data augmentation. We investigated the possibility to train from simulated data with BipartiteEvol, an individual-based model that mimics the eco-evolutionary emergence of bipartite interaction networks (Maliet et al., 2020). However, ANN classifiers trained on BipartiteEvol simulations were not able to discriminate mutualistic and antagonistic empirical networks, suggesting that the model somehow fails at capturing important structural differences between antagonistic and mutualistic networks. (Maliet et al., 2020) showed that the model produces realistic networks, which we confirm here, and is able to reproduce major global differences observed between mutualistic and antagonistic networks. However, we find that BipartiteEvol simulations tend to have higher frequencies of complex motifs and a lower connectance compared to empirical networks. Consequently, their projection only covers a small proportion of the space occupied by empirical networks. We cannot exclude that using different parameters in our simulations would provide more heterogeneous simulated networks that better match the high variability observed in empirical networks. Alternatively, one possible explanation is that many bipartite interaction networks belong to a continuum between antagonism and mutualism rather than a strict dichotomy. Indeed, such a continuum between mutualism and antagonism has been observed in many interactions found in natural communities (Montesinos-Navarro et al., 2017). For instance, it has been shown that parrot-plant networks can combine the two types of interactions (Montesinos-Navarro et al., 2017). Similarly, pollination or mycorrhizal networks often contain cheating species that use antagonistic strategies and induce structural changes within the interaction networks (Genini et al., 2010; Klironomos, 2003; Perez-Lamarque et al., 2020). Interactions can also shift from mutualism to antagonism on short timescales, depending on some external conditions (Canestrari et al., 2014; Sachs et al., 2011). By contrast, BipartiteEvol models the emergence of a network in strict antagonistic or mutualistic communities. This calls for future efforts to improve the BipartiteEvol model or to use simulations under other generative models (*e.g.,* (Cai et al., 2020; Thebault & Fontaine, 2010). Alternatively, data augmentation methods could also be tested to generate synthetic networks by randomly subsampling, mixing and rewiring empirical networks; but whether such approaches produce realistic interaction networks would need to be tested first.

Ultimately, supervised classifying approaches could be particularly useful to predict the type of undetermined interactions. For instance, many plant-fungus interaction networks are assembled from high-throughput sequencing data, but we currently ignore the mutualistic *versus* the antagonistic type of many of these endophytic interactions (Chagnon et al., 2016; Selosse et al., 2018). Similarly, an increasing number of networks are built from co-occurrence data generated with high-throughput sequencings, such as the ones between microscopic planktonic species sampled by the *Tara Oceans* expeditions (Chaffron et al., 2021; Lima-Mendez et al., 2015). The ecology of the majority of species in these networks is currently undetermined (Bjorbækmo et al., 2020; Lima-Mendez et al., 2015; Strydom et al., 2021). However, a preliminary analysis of bipartite networks built from co-occurrence data in the *Tara Oceans* data revealed that these networks were structurally very different from the empirical and simulated networks we used here (results not shown). Classifying these networks would thus require using a specific training dataset composed of microbial networks with known interaction types (as discussed in Supplementary Note 3). Similarly, while our approach is strictly limited to bipartite networks, the increasing availability of multipartite networks, embracing different ecologies (Domínguez-García & Kéfi, 2024; Pocock et al., 2012), and the recent development of generative models for multipartite networks (Hale et al., 2023) suggest that a classification approach to more complex networks may now be possible. Embracing the challenge of telling apart interaction types in multipartite networks from their structure would also allow to evaluate differences in structure between simulated and empirical multipartite networks.

## Conclusion

Mutualistic and antagonistic networks have structural differences that can be particularly well captured by motif frequencies with supervised machine learning. Antagonistic networks are enriched in densely connected motifs, while mutualistic networks are enriched in motifs with asymmetrical specialization. Besides this dichotomy, there are structural differences across networks with distinct ecologies. These structural differences can be used to classify bipartite interaction networks with supervised machine learning with pretty good accuracy. These performances could be further improved by increasing the number of empirical networks used for training, diversifying the type of ecology sampled, and improving currently available models to simulate networks with realistic structures. While more work is needed to predict interaction types for co-occurrence networks, our analyses provide promising results on the ability to predict interaction types from network structure.

The following results were obtained by training the artificial neural networks (ANN) with 10,000 different training and test sets, using motif frequencies as the input structural variables. From those 10,000 classifications, we calculated the percentage of antagonistic *versus* mutualistic classifications for each network: a network was predicted as a mutualistic (resp. antagonistic) network with high confidence (highlighted in grey) if it was classified as a mutualistic (resp. antagonistic) network in more than 75% of the classifications. It was predicted as a mutualistic (resp. antagonistic) network with low confidence if it was classified as a mutualistic (resp. antagonistic) network in more than 50% but less than 75% of the classifications. We therefore obtained a predicted type of interaction for each of the 343 empirical networks. The distributions of these classification confidence for all networks are given in Fig. S18.

The second column indicates the number of interaction networks in our database gathered for each ecology of interaction. Then, we indicate the number (and the percentage) of networks classified as either mutualistic or antagonistic for each ecology of interaction and at the bottom, we show the global contingency table for all mutualistic and antagonistic networks. We also indicate the F-score associated with the global contingency table for each antagonistic or mutualistic type of interaction. Rows are ordered based on the % of networks classified as mutualistic within each type of ecology. In the end, 88 antagonistic and 191 mutualistic networks are correctly classified, i.e., 279 of the 343 empirical networks (81%).

## Acknowledgments

We thank I. Overcast, J. Voznica, L. Aristide, S. Lambert, I. Quintero, C. Fruciano, J. Clavel, A. Silva, for comments on the first version of the manuscript. We also thank S. Chaffron, C. Bowler, C. Nef, and E. Thébault for fruitful discussions. We thank the three reviewers for their comments that improved the manuscript.

The authors declare no conflict of interest.

## Author contributions

All authors conceived the ideas and designed methodology; B.P performed research. B.P, H.M and B.P.L led the writing of the manuscript. All authors contributed critically to the drafts and gave final approval for publication.

## Data availability

The code used for the analyses is available on Zenodo: https://doi.org/10.5281/zenodo.7786771

