## supplementary for "Telling mutualistic and antagonistic ecological networks apart by learning their multiscale structure"

**Supplementary Notes (1-3)**

**Supplementary Tables (1-9)**

**Supplementary Figures (1-18)**

### Supplementary Notes:

##### Supplementary Note 1: Using simulated networks to increase the number of networks in the training set

We simulated the BipartiteEvol model using a wide range of antagonistic and mutualistic parameters (see Perez-Lamarque, Maliet, et al., 2022, for details) and obtained 1,200 weighted networks. We noticed that connectance values for these networks were on average lower than those of the empirical networks; to approach those of the empirical networks, we retained 70% randomly sampled individual interactions in the simulated networks, which had a direct effect of deleting rare, specialized species and increasing network connectance. We also carried out analyses with the original networks (no sampling), which did not qualitatively affect the results (results not shown). Retaining networks that had at least 10 species in each guild, we obtained a total of 1,175 binary networks, including 705 mutualistic and 470 antagonistic networks, hereafter referred to as BipartiteEvol networks.

We performed unsupervised classification using PCA and K-means. We also performed supervised classification: we independently trained and tested the supervised classifiers (Logit or Lasso regressions, or ANN) on BipartiteEvol networks using the different sets of structural input variables. We then tested the classifier on the empirical networks. We investigated differences between networks simulated with BipartiteEvol and empirical networks by comparing the distributions of their global metrics (in terms of network size, connectance, nestedness, and modularity), their motifs frequencies, and the spectral densities of their Laplacian graph.

#### Supplementary Note 2: Differences in network structure between empirical networks and BipartiteEvol simulated networks

To understand why the simulations failed at improving the classification accuracy of empirical networks, we compared one-by-one the global metrics of the simulated and the empirical networks. We found networks simulated using BipartiteEvol to be on average significantly more nested and less modular than empirical ones (Fig. S10). They also remained slightly less connected (0.11) than empirical networks (0.19) despite the strategy we used to increase their connectance. Principal component analyses (PCA) using either global metrics, motifs frequencies, or spectral density values indicated that BipartiteEvol networks are clustered in a limited space within the space occupied by the empirical networks. This indicates that our simulations are realistic but mimic only a fraction of what empirical networks can look like (Figs. S11-13). In addition, BipartiteEvol networks often had significantly higher frequencies of complex motifs (involving 6 nodes and highly connected) compared with empirical networks that were often enriched with simpler motifs (Fig. S14). In line with these results, the Laplacian density of BipartiteEvol networks was characterized by larger eigenvalues than those of empirical networks (Fig. S15). Altogether, networks simulated using BipartiteEvol are not as structurally diverse as empirical networks and slightly distinct in their structure, explaining at least partly why ANN trained on BipartiteEvol networks performed badly at classifying empirical networks (Table S8B).

#### Supplementary Note 3: Discussing the use of ANN to predict the type of interaction in networks built from OTU

Preleminary analysis on bipartite networks built from co-occurrence date in the Tara Oceans data revealed that our classification approach failed at classifying networks with known type of interactions. A potential source of difference might come from species delineation in microbial lineages in such databases, which rely on the clustering into operational taxonomic units (OTUs) of short marker gene sequences (*e.g.* the small subunit rRNA gene or the ITS marker). This difference in species delimitation in networks involving microbial groups compared to networks involving macro-organisms (*e.g.* pollination and herbivory) (Chagnon, 2016) may affect network structure. This may explain why the classification of mycorrhizal networks was not very accurate in our study. One approach to tackle this would be to incorporate more microbial networks with known interaction types in the training of the supervised classifier, which would be highly recommended anyway given the importance of the type of ecology in the classification accuracy. Another source of differences comes from the use of co-occurrence as a proxy for interaction, a topic that is currently intensively debated (Blanchet et al., 2020). Hence, we suggest that to apply our approach to microbial groups, one would need to train the ANN with co-occurrence networks with known interaction types or simulated co-occurrence networks.

### Supplementary Tables:

**Supplementary Table 1: Differences in global metrics are better explained by the type of ecology than by the antagonistic/mutualistic type of interactions.**

We fitted linear and mixed-effects models to compare whether the type of ecology (e.g. pollination, herbivory…) or the type of interactions (i.e. mutualistic *versus* antagonistic) better explained the variation in the global metrics (nestedness, modularity, connectance, and both difference and sum of guild size). In the first column, separate linear mixed models were performed for each global metric with the type of interactions as a fixed effect and the type of ecology as a random one. The second column corresponds to a linear model for each global metric with the type of ecology as the only fixed effect. And in the last column, we performed linear models with the type of interactions as the only fixed effect.

For each model, we reported the model fit, using the Akaike information criterion (AIC). The lower the AIC, the better the fit, and for each global metric, the best model is indicated in bold.

| | **AIC: antagonistic/mutualistic (fixed effect) accounting for the type of ecology (random effect)** | **AIC: type of ecology (fixed effect)** | **AIC: antagonistic/mutualistic (fixed effect)** |
|---|---|---|---|
| Nestedness | 819.6300 | **801.3235** | 879.6975 |
| Connectance | -545.3387 | **-575.7837** | -462.6151 |
| Modularity | -472.6396 | **-494.5077** | -467.7184 |
| Difference of guild size | **3633.3023** | 3638.1984 | 3644.7096 |
| Sum of guild size | **3977.3292** | 3987.1752 | 3988.2689 |

**Supplementary Table 2: Difference in global metrics between the different types of ecology in empirical networks.**

We compared the differences in global metrics between the different types of ecology by performing a separate ANOVA followed by post hoc Tukey tests on the connectance (A), modularity (B), and nestedness (C). We used the Bonferroni adjustment to account for multiple tests. Here, we compared the difference in global metrics between each pair of networks from different types of ecology. We excluded from this analysis anemone-fish and ant-plant as they were represented by a small number of networks (1 and 3 respectively).

**A**

| Types of ecology compared | Mean difference | P-value |
|---|---|---|
| Herbivory-Bacteria-Phage | **-0.34** | <0.001 |
| Host-Parasite-Bacteria-Phage | **-0.25** | <0.001 |
| Host-Parasitoid-Bacteria-Phage | **-0.36** | <0.001 |
| Mycorrhiza-Bacteria-Phage | **-0.32** | <0.001 |
| Pollination-Bacteria-Phage | **-0.33** | <0.001 |
| Seed dispersal-Bacteria-Phage | **-0.25** | <0.001 |
| Host-Parasite-Herbivory | **0.10** | 0.002 |
| Host-Parasitoid-Herbivory | -0.01 | 1.000 |
| Mycorrhiza-Herbivory | 0.02 | 0.998 |
| Pollination-Herbivory | 0.01 | 0.999 |
| Seed dispersal-Herbivory | **0.10** | 0.013 |
| Host-Parasitoid-Host-Parasite | **-0.11** | 0.027 |
| Mycorrhiza-Host-Parasite | -0.07 | 0.389 |
| Pollination-Host-Parasite | **-0.09** | <0.001 |
| Seed dispersal-Host-Parasite | 0.00 | 1.000 |
| Mycorrhiza-Host-Parasitoid | 0.04 | 0.989 |
| Pollination-Host-Parasitoid | 0.02 | 0.991 |
| Seed dispersal-Host-Parasitoid | 0.11 | 0.052 |
| Pollination-Mycorrhiza | -0.01 | 1.000 |
| Herbivory-Bacteria-Phage | -0.34 | 0.466 |
| Host-Parasite-Bacteria-Phage | **-0.25** | 0.001 |

**B**

| Types of ecology compared | Mean difference | P-value |
|---|---|---|
| Herbivory-Bacteria-Phage | 0.07 | 0.567 |
| Host-Parasite-Bacteria-Phage | -0.06 | 0.368 |
| Host-Parasitoid-Bacteria-Phage | 0.09 | 0.438 |
| Mycorrhiza-Bacteria-Phage | 0.03 | 0.991 |
| Pollination-Bacteria-Phage | 0.00 | 1.000 |
| Seed dispersal-Bacteria-Phage | -0.02 | 0.998 |
| Host-Parasite-Herbivory | **-0.13** | <0.001 |
| Host-Parasitoid-Herbivory | 0.03 | 0.998 |
| Mycorrhiza-Herbivory | -0.03 | 0.994 |
| Pollination-Herbivory | -0.06 | 0.189 |
| Seed dispersal-Herbivory | -0.09 | 0.116 |
| Host-Parasitoid-Host-Parasite | **0.16** | 0.002 |
| Mycorrhiza-Host-Parasite | 0.10 | 0.187 |
| Pollination-Host-Parasite | **0.07** | <0.001 |
| Seed dispersal-Host-Parasite | 0.04 | 0.569 |
| Mycorrhiza-Host-Parasitoid | -0.06 | 0.941 |
| Pollination-Host-Parasitoid | -0.09 | 0.236 |
| Seed dispersal-Host-Parasitoid | -0.11 | 0.128 |
| Pollination-Mycorrhiza | -0.03 | 0.985 |
| Seed dispersal-Mycorrhiza | -0.05 | 0.879 |
| Seed dispersal-Pollination | -0.02 | 0.955 |

C

| Types of ecology compared | Mean difference | P-value |
|---|---|---|
| Herbivory-Bacteria-Phage | **1.98** | <0.001 |
| Host-Parasite-Bacteria-Phage | 0.45 | 0.295 |
| Host-Parasitoid-Bacteria-Phage | **1.56** | <0.001 |
| Mycorrhiza-Bacteria-Phage | 0.62 | 0.434 |
| Pollination-Bacteria-Phage | **1.02** | <0.001 |
| Seed dispersal-Bacteria-Phage | **0.81** | 0.011 |
| Host-Parasite-Herbivory | **-1.53** | <0.001 |
| Host-Parasitoid-Herbivory | -0.42 | 0.780 |
| Mycorrhiza-Herbivory | **-1.36** | <0.001 |
| Pollination-Herbivory | **-0.96** | <0.001 |
| Seed dispersal-Herbivory | **-1.17** | <0.001 |
| Host-Parasitoid-Host-Parasite | **1.11** | <0.001 |
| Mycorrhiza-Host-Parasite | 0.17 | 0.996 |
| Pollination-Host-Parasite | **0.57** | <0.001 |
| Seed dispersal-Host-Parasite | 0.36 | 0.335 |
| Mycorrhiza-Host-Parasitoid | -0.94 | 0.109 |
| Pollination-Host-Parasitoid | -0.54 | 0.308 |
| Seed dispersal-Host-Parasitoid | -0.75 | 0.105 |
| Pollination-Mycorrhiza | 0.40 | 0.735 |
| Herbivory-Bacteria-Phage | 0.19 | 0.996 |
| Host-Parasite-Bacteria-Phage | -0.21 | 0.816 |

**Supplementary Table 3: K-means on PCA projections failed at discriminating networks with different types of ecology.**

K-means classification using the two principal components analysis of the PCA performed using global metrics (nestedness, modularity, connectance, and the two metrics of network sizes) in panel A, spectral densities of the Laplacian matrix in panel B, and motifs frequencies in panel C. We indicate the Chi-squared test. In addition, for each type of ecology, we also indicate the percentage of networks classified outside the cluster containing most of the networks of this type of ecology. For instance, using Kmeans on PCA projection based on global metrics led to 21 herbivory networks in cluster 1 and 2 in cluster 2. We therefore indicate the percentage 2/23 = 8.7% to emphasize that a majority of herbivory networks are classified in the same Kmeans cluster. df=NA because we used a Monte Carlo approach to compute the $p$-values.

### A = Global metrics

| | Anemone-Fish | Bacteria-Phage | Herbivory | Host-Parasite | Host-Parasitoid | Mycorrhiza | Ant-Plant | Pollination | Seed dispersal |
|---|---|---|---|---|---|---|---|---|---|
| Cluster 1 | 1 | 17 | 21 | 70 | 10 | 7 | 3 | 162 | 27 |
| Cluster 2 | 0 | 0 | 2 | 7 | 0 | 2 | 0 | 12 | 2 |
| Chi² test | X²= 237.19, df = NA, p-value <0.001 | | | | | | | | |
| % of networks classified outside the main cluster for each type of ecology | 0 | 0 | 8.7 | 9.1 | 0 | 22.2 | 0 | 6.9 | 6.9 |

### B = Spectral density of the Laplacian graph

| | Anemone-Fish | Bacteria-Phage | Herbivory | Host-Parasite | Host-Parasitoid | Mycorrhiza | Ant-Plant | Pollination | Seed dispersal |
|---|---|---|---|---|---|---|---|---|---|
| Cluster 1 | 0 | 6 | 19 | 40 | 10 | 7 | 2 | 159 | 11 |
| Cluster 2 | 1 | 11 | 4 | 37 | 0 | 2 | 1 | 15 | 18 |
| Chi² test | X²= 205.3, df = NA, p-value <0.001 | | | | | | | | |
| % of networks classified outside the main cluster for each type of ecology | 0 | 35.3 | 17.4 | 48.1 | 0 | 22.2 | 33.3 | 8.6 | 37.9 |

### C = Motif frequencies

| | Anemone-Fish | Bacteria-Phage | Herbivory | Host-Parasite | Host-Parasitoid | Mycorrhiza | Ant-Plant | Pollination | Seed dispersal |
|---|---|---|---|---|---|---|---|---|---|
| Cluster 1 | 0 | 14 | 1 | 34 | 0 | 1 | 1 | 5 | 6 |
| Cluster 2 | 1 | 3 | 22 | 43 | 10 | 8 | 2 | 169 | 23 |
| Chi² test | X² = 330.34, df = NA, p-value <0.001 | | | | | | | | |
| % of networks classified outside the main cluster for each type of ecology | 0 | 17.6 | 4.3 | 44.2 | 0 | 11.1 | 33.3 | 2.9 | 20.7 |

**Supplementary Table 4: Classification accuracy of the empirical networks using artificial neural networks (ANN), Lasso regressions, or logit regressions**.

Each supervised clustering method was tested with the different input variables used to characterize network structure (global metrics, motif frequencies, and spectral density values), as well as any combination of these sets of structural variables. For each combination of input variables and methods, we used 50 independent training and testing sets. The mean of the percentages of correct classifications are indicated for antagonistic (yellow) and mutualistic (green) networks. The F-score is indicated in parenthesis.

| | global metrics | | motifs | | spectral density | | global metrics + spectral density | | global metrics + motifs | | spectral density + motifs | | spectral density + motifs + global metrics | |
|---|---|---|---|---|---|---|---|---|---|---|---|---|---|---|
| **Logit** | 31 (0.43) | 93 (0.8) | 58 (0.63) | 84 (0.81) | 55 (0.53) | 70 (0.71) | 58 (0.56) | 71 (0.73) | 63 (0.66) | 84 (0.82) | 62 (0.59) | 71 (0.73) | 64 (0.6) | 72 (0.75) |
| **Lasso** | 26 (0.39) | 95 (0.8) | 49 (0.61) | 93 (0.83) | 39 (0.49) | 89 (0.79) | 42 (0.53) | 90 (0.8) | 52 (0.63) | 92 (0.84) | 53 (0.64) | 92 (0.84) | 52 (0.62) | 90 (0.82) |
| **Neural network** | 53 (0.61) | 88 (0.82) | 65 (0.68) | 84 (0.82) | 57 (0.59) | 79 (0.77) | 58 (0.6) | 79 (0.78) | 65 (0.66) | 82 (0.81) | 65 (0.66) | 82 (0.81) | 60 (0.62) | 81 (0.79) |
| | *Antagonistic networks*<br>*Mutualistic networks* | | | | | | | | | | | | | |

**Supplementary Table 5: Influence of the bandwidth used to transform the Laplacian spectrum into a density on the classification of the empirical networks.**

We compared the percentages of correct classifications of the empirical networks based on artificial neural network (ANN) using the spectral density of the Laplacian graph for different bandwidths (from 0.001 to 0.091) as the inputs. High bandwidth values tend to smooth the densities (irrespective of the network size), whereas low bandwidth values tend to increase the details in the densities and are thus more sensitive to the network size. Training and testing were repeated 50 times of different training and test sets, and the table represents the mean percentage of correct classifications.

* 0.025 is the value used in all other analyses.

| Bandwidth | Percentage of correct classifications for antagonistic networks | Percentage of correct classifications for mutualistic networks |
|---|---|---|
| **0.001** | 58 % | 70 % |
| **0.011** | 60 % | 78 % |
| **0.021** | 64 % | 78 % |
| **0.025 *** | 65 % | 76 % |
| **0.031** | 63 % | 76 % |
| **0.041** | 61 % | 75 % |
| **0.051** | 65 % | 77 % |
| **0.061** | 64 % | 74 % |
| **0.071** | 65 % | 73 % |
| **0.081** | 65 % | 75 % |
| **0.091** | 65 % | 73 % |

**Supplementary Table 6: Low overfitting in the Lasso and Probit classification using empirical network motifs.**

We trained Probit and Lasso methods on motif frequencies of empirical networks and tested them on the same set of networks (the training set). All numbers in the tables correspond to the mean percentages of classifications in each interaction type for 50 different training sets.

| Probit | **Testing on the training set** | |
|---|---|---|
| | Antagonistic | Mutualistic |
| Antagonistic | 72 | 28 |
| Mutualistic | 8 | 92 |

| Lasso | **Testing on the training set** | |
|---|---|---|
| | Antagonistic | Mutualistic |
| Antagonistic | 54 | 46 |
| Mutualistic | 5 | 95 |

**Supplementary Table 7: ANN classification is influenced by the signal from each type of ecology.**

We excluded one by one a type of ecology from the training test (e.g. pollination networks), we trained the ANN (using motifs frequencies as inputs) on 50 independent training sets constituted by 80% of the remaining networks, and tested the classification on the networks from the removed type of ecology. All numbers in the tables correspond to the mean percentage of correct classifications and the associated standard deviation.

Percentage of correct classifications of the networks belonging to the type of ecology removed from the training set. In the right column, we also display the original percentage of correct classifications when all the networks are used for comparison. This percentage is in general higher, indicating that the network classification improves when networks of the same ecology are used in the training. However, the networks tend to be properly classified even if networks of their ecology were not included in the training sets, indicating that the classifier learns the antagonist *versus* mutualist interaction types besides the ecology. Exceptions are the host-parasite and pollination networks (*), which represent 60% and 80% of the antagonist and mutualist networks, respectively, and which simply do not let enough representatives of their interaction type once removed. The herbivory networks (**) are also poorly classified, but it is the case even when they are included in the training set.

| **Type of ecology excluded** | **Percentage of correct classifications for the networks excluded from the training set** | **Percentage of correct classifications with all the networks in the training set** |
|---|---|---|
| Bacteria-Phage | 58 % +/- 11.9 | 71 % |
| Seed dispersal | 65 % +/- 5.6 | 79 % |
| Host-Parasite* | 25 % +/- 5.3 | 78 % |
| Host-Parasitoid | 51 % +/- 8.2 | 50 % |
| Mycorrhiza | 60 % +/- 15.1 | 55 % |
| Plant-Ant | 63 % +/- 22.9 | 66 % |
| Herbivory** | 18 % +/- 8.5 | 43 % |
| Pollination* | 26 % +/- 4.6 | 92 % |

**Supplementary Table 8: BipartiteEvol simulations failed at classifying antagonistic and mutualistic networks based on their structures, even simulated ones.**

Results from artificial neural networks (ANN) trained to classify antagonistic *versus* mutualistic networks using motif frequencies as input variables. The training was repeated 50 times with different training sets constituted by 80% of the BipartiteEvol simulated networks. Then, each classifier was tested on the 20% remaining simulated networks (A) or on all the empirical networks (B). The mean percentages of classifications is indicated for antagonistic (yellow) and mutualistic ones (green).

For each classification, the F-score is also indicated.

A) Tests on the 20% remaining simulated networks

| | Antagonistic | Mutualistic |
|---|---|---|
| Antagonistic | 70 | 30 |
| Mutualistic | 42 | 58 |
| F-scores | 0.71 | 0.57 |

B) Tests on the empirical networks

| | Antagonistic | Mutualistic |
|---|---|---|
| Antagonistic | 71 | 29 |
| Mutualistic | 86 | 14 |
| F-scores | 0.45 | 0.21 |

**Supplementary Table 9: Classification accuracy increases with the complexity of the motifs involved.**

Results of Artificial neural networks (ANN) classifications of antagonistic *versus* mutualistic networks using frequencies of motifs of size at most 4, 5 and 6 (all motifs) as inputs. For classifications using at most 4- (*resp.* 5-) nodes motifs, we obtained each motif frequency by dividing the number of each of these motifs by the total number of motifs involved. The training was repeated 50 times with different training sets constituted of 80% of all empirical networks. Then, each classifier was tested on the 20% remaining simulated networks. The mean of the percentages of correct classifications and F-scores (in parenthesis) are indicated for either antagonistic networks (yellow) or mutualistic ones (green).

| Motifs used | **% of correct classification of mutualistic networks** | **% of correct classification of antagonistic networks** |
|---|---|---|
| All motifs with at most 4-nodes | 80% (F-score=0.79) | 63% (F-score=0.65) |
| All motifs with at most 5-nodes | 82% (F-score=0.81) | 67% (F-score=0.68) |
| All motifs (Table S4) | 65% (F-score=0.68) | 84% (F-score=0.82) |

### Supplementary Figures:

**Supplementary Figure 1: Global metrics of the empirical networks vary according to the different types of ecology.**

The connectance (A), nestedness (B; NODFc), modularity (C; Newman's index), the difference (D), and sum (E) of guilds size of the empirical networks are represented for each type of ecology (pollination, herbivory…). The mutualistic networks are colored in green and the antagonistic ones, in yellow.

Boxplots present the median surrounded by the first and third quartiles, and whiskers extend to the extreme values but no further than 1.5 of the interquartile range. For visualization issues we limited the values of sizes to 100 (resp. 200) for panel D (resp. E), therefore omitting 22 networks in both cases.

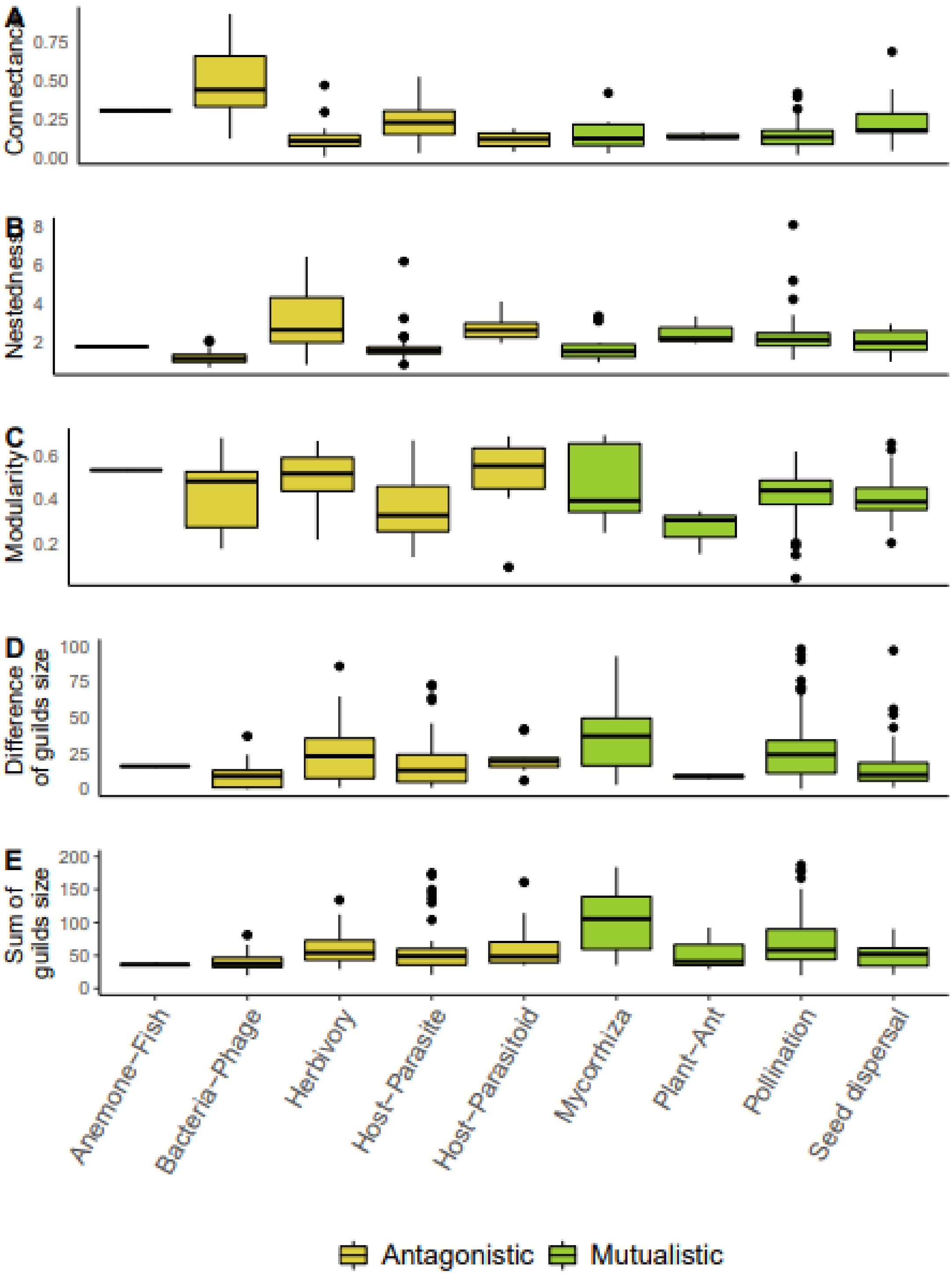
A
Connectance
0.00
0.25
0.50
0.75
B
Nestedness
2
4
6
8
C
Modularity
0.2
0.4
0.6
D
Difference of guilds size
0
25
50
75
100
E
Sum of guilds size
0
50
100
150
200
Anemone−Fish
Bacteria−Phage
Herbivory
Host−Parasite
Host−Parasitoid
Mycorrhiza
Plant−Ant
Pollination
Seed dispersal
Antagonistic
Mutualistic

**Supplementary Figure 2: The Laplacian spectral density, a multi-scale approach to study the structure of ecological networks**

(A) Illustration of the computation of the spectral density of the Laplacian graph associated with an interaction network. From the matrix depicting species interactions (here, the ant-plant interaction network from Passmore *et al.* 2012), we compute the normalized Laplacian graph ($L_{norm}$) and its associated eigenvalues (we note $\lambda_i$ the i-th eigenvalue and N the number of eigenvalues). The spectral density $f(x)$ represents the density of eigenvalues, obtained using a gaussian kernel of parameter $\sigma$, fixed here at 0.025.

(B-E) Four examples of spectral densities computed on a perfectly modular network (B), a perfectly nested network (C), a host-parasite network (D; Bellay *et al.* 2013), and a pollination network (E; Gilarranz *et al.* 2015). Modularity (M; Newman's modularity), nestedness (N; NODFc) and the size of the guilds are indicated for each network. For presentation purposes, the empirical networks were not represented with their true dimensions.

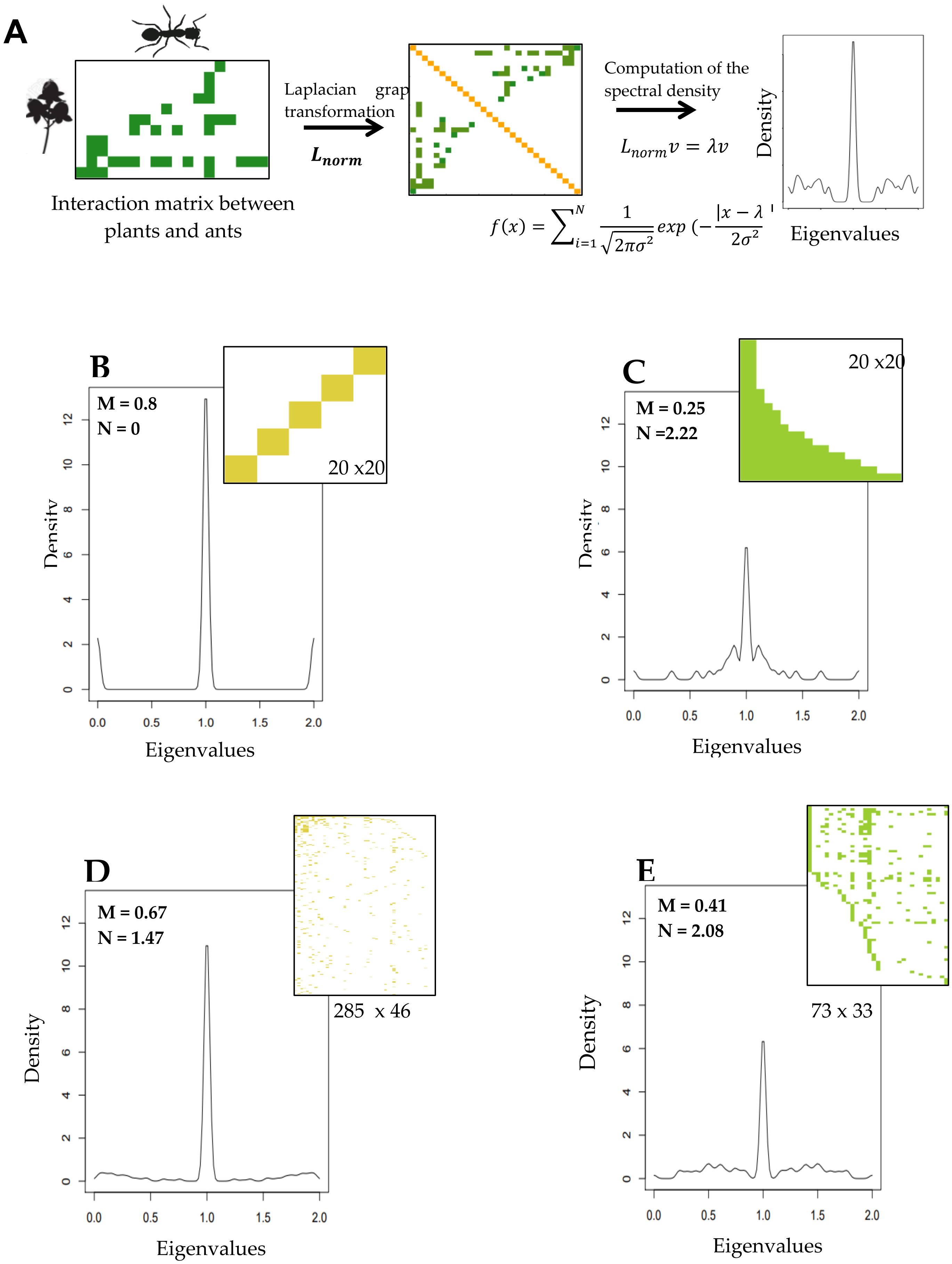
A
Interaction matrix between plants and ants
Laplacian graph transformation
$L_{norm}$
Computation of the spectral density
$L_{norm}v = \lambda v$
$f(x) = \sum_{i=1}^{N} \frac{1}{\sqrt{2\pi\sigma^2}} exp\,(-\frac{|x-\lambda|}{2\sigma^2}$
Density
Eigenvalues
B
M = 0.8
N = 0
20 x20
C
M = 0.25
N =2.22
20 x20
D
M = 0.67
N = 1.47
285 x 46
E
M = 0.41
N = 2.08
73 x 33

**Supplementary Figure 3: The type of ecology influences the shape of the spectral density of the Laplacian graph**

We compared the spectral density of the Laplacian matrix between the different types of ecology in empirical networks (A: 0 to 0.25; B: 0.25 to 0.5; C: 0.5 to 0.75; D: 0.75 to 1). For that, we used linear models for each density of eigenvalue with the type of ecology as the only effect (Fisher test: p-value<0.01); for visualization issues, we only kept the type of ecology for which the number of available networks is higher than 20. Boxplots present the median surrounded by the first and third quartiles, and whiskers extend to the extreme values but no further than 1.5 of the interquartile range.

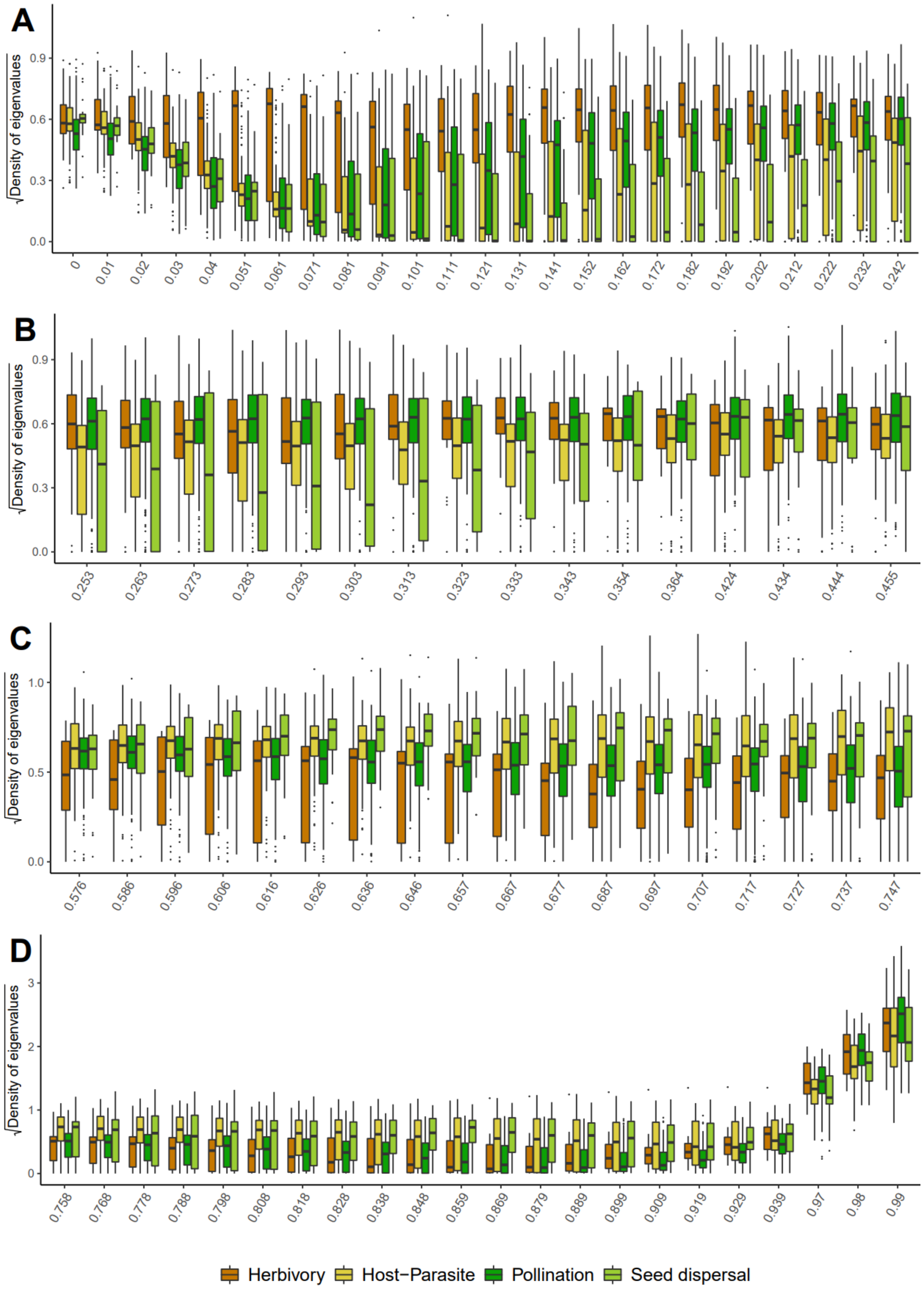
A
B
C
D
√Density of eigenvalues
0 0.01 0.02 0.03 0.04 0.051 0.061 0.071 0.081 0.091 0.101 0.111 0.121 0.131 0.141 0.152 0.162 0.172 0.182 0.192 0.202 0.212 0.222 0.232 0.242
0.253 0.263 0.273 0.283 0.293 0.303 0.313 0.323 0.333 0.343 0.354 0.364 0.424 0.434 0.444 0.455
0.576 0.586 0.596 0.606 0.616 0.626 0.636 0.646 0.657 0.667 0.677 0.687 0.697 0.707 0.717 0.727 0.737 0.747
0.758 0.768 0.778 0.788 0.798 0.808 0.818 0.828 0.838 0.848 0.859 0.869 0.879 0.889 0.899 0.909 0.919 0.929 0.939 0.97 0.98 0.99
Herbivory
Host−Parasite
Pollination
Seed dispersal

**Supplementary Figure 4: Motif frequencies that differ significantly across empirical networks.**

We represented here the motif frequencies that differ significantly when considering the type of interactions alone (A), the type of interactions corrected by the type of ecology (mixed model) (B), and the type of ecology alone (C). In panel A, motifs were selected if their frequency differed significantly between antagonist and mutualist networks (Wilcoxon-Mann-Whitney test: p-value<0.01). In panel B, separate mixed models were performed on each motif with the type of interactions as a fixed effect and the type of ecology as a random effect (Wald test: p-value <0.01). In panel C, linear models were used for each motif with the type of ecology as the only effect (Fisher test: p-value<0.01); for visualization purposes, we only kept the types of ecology that were represented by at least 20 networks. Boxplots present the median surrounded by the first and third quartiles, and whiskers extend to the extreme values but no further than 1.5 of the interquartile range.

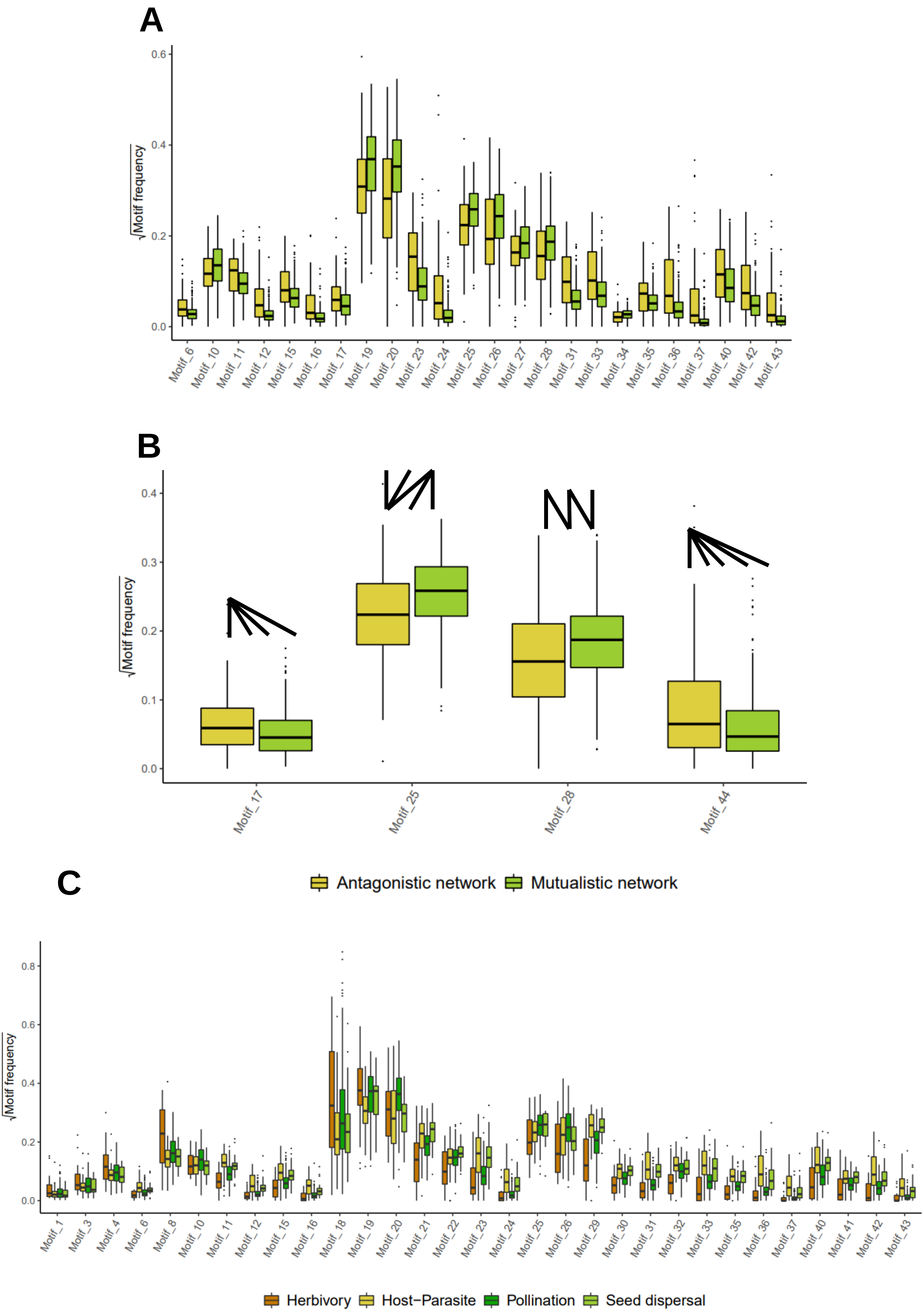
A
√Motif frequency
0.0
0.2
0.4
0.6
Motif_6
Motif_10
Motif_11
Motif_12
Motif_15
Motif_16
Motif_17
Motif_19
Motif_20
Motif_23
Motif_24
Motif_25
Motif_26
Motif_27
Motif_28
Motif_31
Motif_33
Motif_34
Motif_35
Motif_36
Motif_37
Motif_40
Motif_42
Motif_43
B
√Motif frequency
0.0
0.1
0.2
0.3
0.4
Motif_17
Motif_25
Motif_28
Motif_44
Antagonistic network
Mutualistic network
C
√Motif frequency
0.0
0.2
0.4
0.6
0.8
Motif_1
Motif_3
Motif_4
Motif_6
Motif_8
Motif_10
Motif_11
Motif_12
Motif_15
Motif_16
Motif_18
Motif_19
Motif_20
Motif_21
Motif_22
Motif_23
Motif_24
Motif_25
Motif_26
Motif_29
Motif_30
Motif_31
Motif_32
Motif_33
Motif_35
Motif_36
Motif_37
Motif_40
Motif_41
Motif_42
Motif_43
Herbivory
Host−Parasite
Pollination
Seed dispersal

**Supplementary Figure 5: Networks with different types of ecology do not form distinct clusters in a Principal coordinate analysis (PCA) on the global metrics.**

PCA were performed using global metrics (nestedness, modularity, connectance, and the two metrics of network sizes) and we represented the projection on the two principal components (PC1 and PC2) of the empirical networks by type of ecology. The percentage of explained variance is respectively 43.6% and 29.8% for the two first principal components of the PCA.

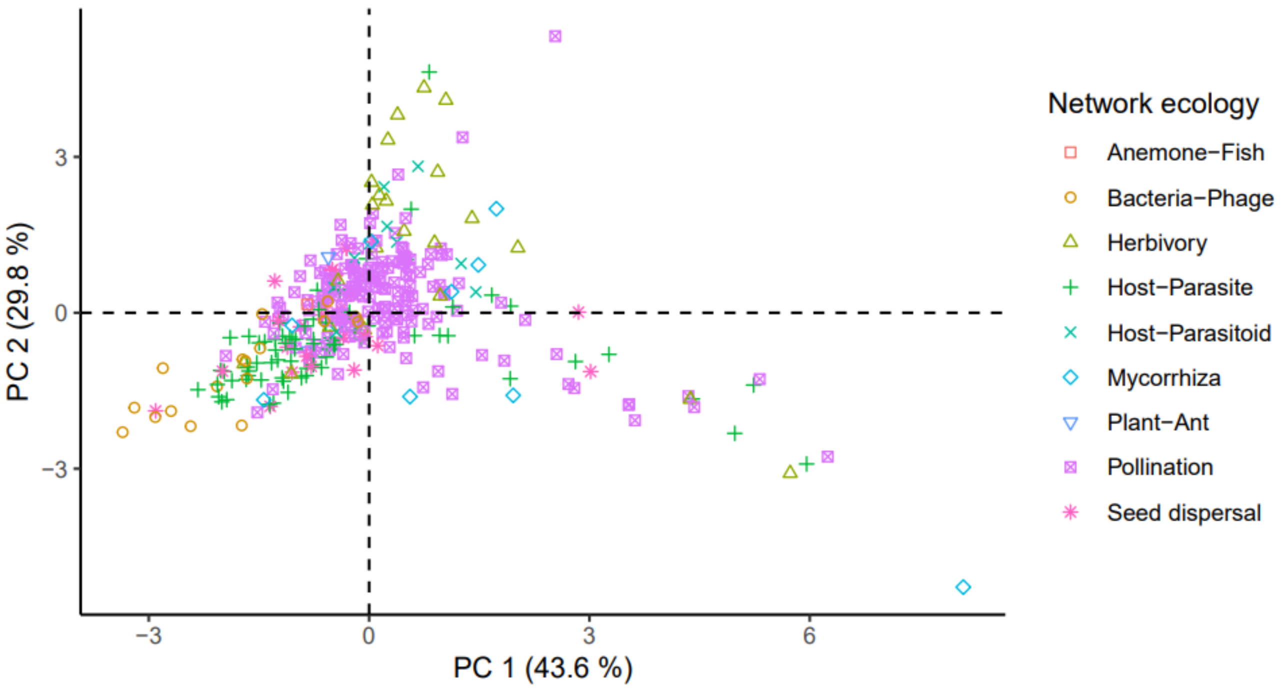

**Supplementary Figure 6: Networks with different types of ecology do not form distinct clusters in a principal coordinate analysis (PCA) on the Laplacian spectral densities.**

PCA was performed using the Laplacian spectral density values and we represented the projection on the two principal components (PC1 and PC2) of the empirical networks by type of ecology. The percentage of explained variance is respectively 31.8 % and 13.9 % for the two first principal components of the PCA.

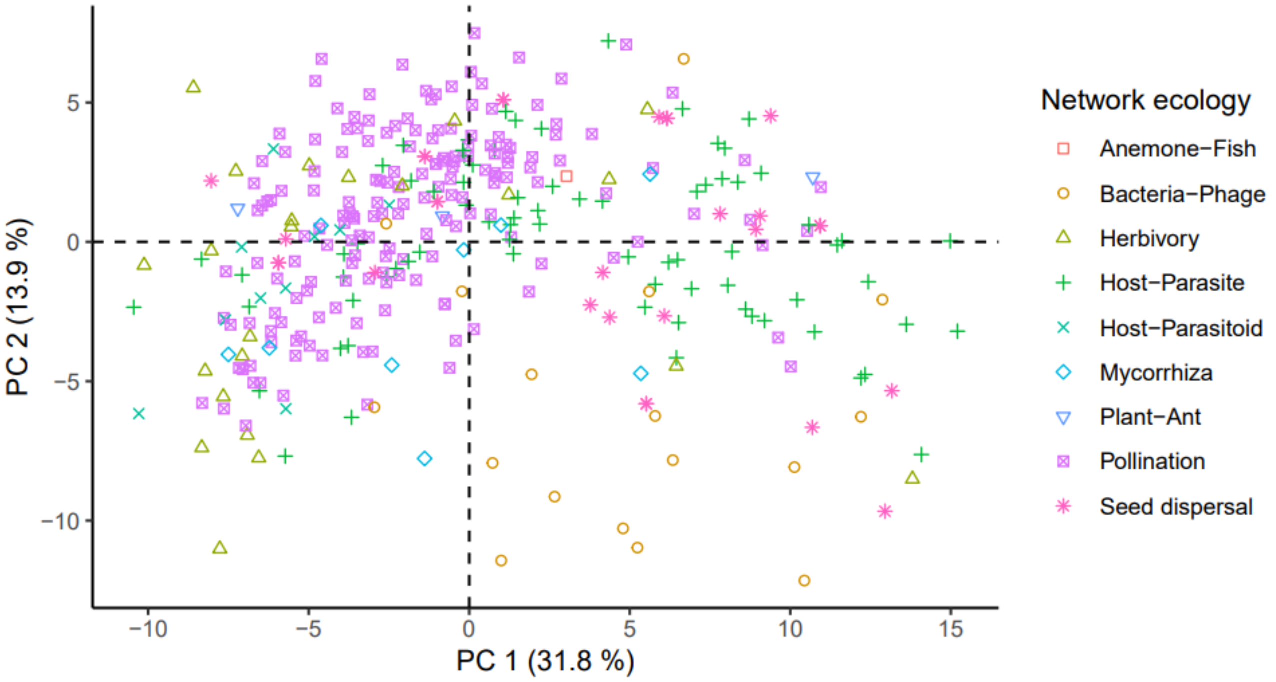

**Supplementary Figure 7: The principal coordinate analysis (PCA) performed on the motif frequencies of the empirical networks tends to discriminate some type of ecology.**

PCA was performed using motif frequencies and we represented the projection on the two principal components (PC1 and PC2) of both empirical and simulated networks. The percentage of explained variance is respectively 28.6 % and 21 % for the two first principal components of the PCA.

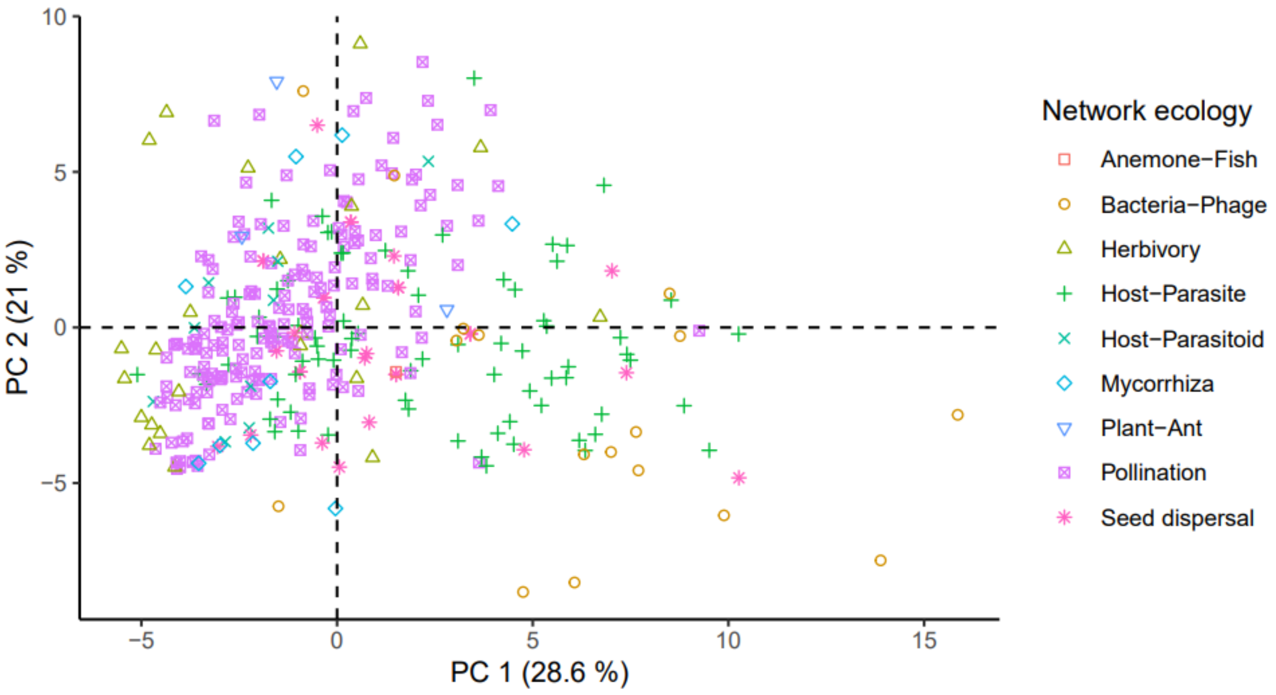

**Supplementary Figure 8: Low effect of overfitting on the percentage of correct classifications of the artificial neural networks (ANNs) trained with the motif frequencies of the empirical networks.**

We trained ANNs with different hidden layer sizes (number of neurons in the intermediate layer growing from 0 to 40) and tested it either with the same set of networks (training set) or with the remaining networks (test set) using motif frequencies as input variables.

The black dot corresponds to the percentages of correct classifications of the method on the training set, whereas the green and yellow dots correspond to the percentages of correct classifications of the method on the test sets for mutualistic and antagonistic networks respectively. Black dots higher than the green/yellow dots indicate overfitting. However, the percentages of correct classifications on the test sets (green/yellow lines) is not strongly affected by overfitting.

All numbers correspond to the mean for 50 different training and test sets.

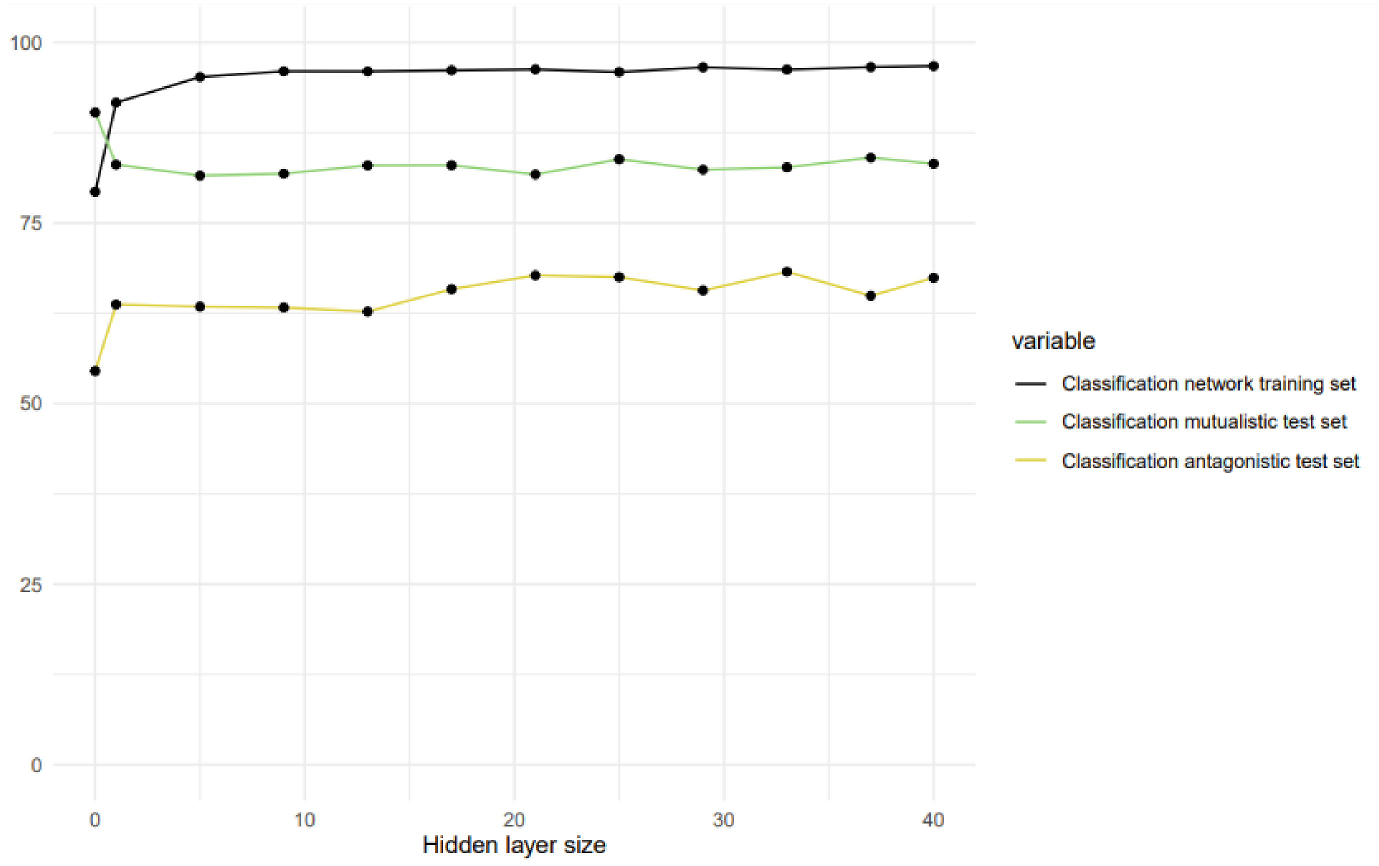

**Supplementary Figure 9: The heterogeneity in the number of networks per type of interaction (A) and type of ecology (B) in the training set only moderately affects the accuracy of the classification.**

(A) We trained the ANN with a training set having the same number (from 5 to 95) of mutualistic and antagonistic networks (“controlled”; dashed lines) and tested it on the test set (*i.e.* remaining networks). We compared the resulting percentage of correct classifications to the one obtained when the ANN is trained with a training set of the same size but with randomly chosen networks (i.e. same total number of networks, but without controlling per type of interaction; continuous lines). Each point corresponds to the mean of 50 independent training and test sets. Controlling for the asymmetry in the number of representants of each type of interaction in the training set leads to a more symmetrical classification accuracy between the two types: it increases by 9.8 % (*resp*. decreases by 8.8%) the percentag of correct classifications of antagonistic (*resp.* mutualistic) networks. On average, the percentage of correct classifications is similar as when not controlling for the asymmetry.

(B) We trained the ANN while constraining the number of networks per type of ecology in the training set to a maximal value (“controlled”; dashed lines; from 5 to 20) and tested it on the test set (*i.e.* remaining networks). We compared this classification with the one where ANN is trained with a training set of the same size but without the constraint on the type of ecology (*i.e.* randomly chosen networks; continuous lines). We indicated at the top of the plot the number of networks in the training set for each point. Each point corresponds to the mean of 50 training and test sets. Similar to above, controlling for the heterogeneity in the number of networks per type of ecology increases the symmetry of classification accuracy between the two interaction types and leads to similar overall percentage of correct classifications on average.

In both cases, when we control for an equal number of networks per (A) type of interaction and (B) type of ecology in the training set, we increase the percentage of correct classifications of antagonistic networks while decreasing the percentage of correct classifications of mutualistic networks. When mutualistic networks are over-represented in the training set, networks that have a rather non-specific structure with respect to interaction type tend to be classified as mutualistic; this does not occur when the training set is more balanced. In all cases, the average percentage of correct classifications is good (around 75%).

A

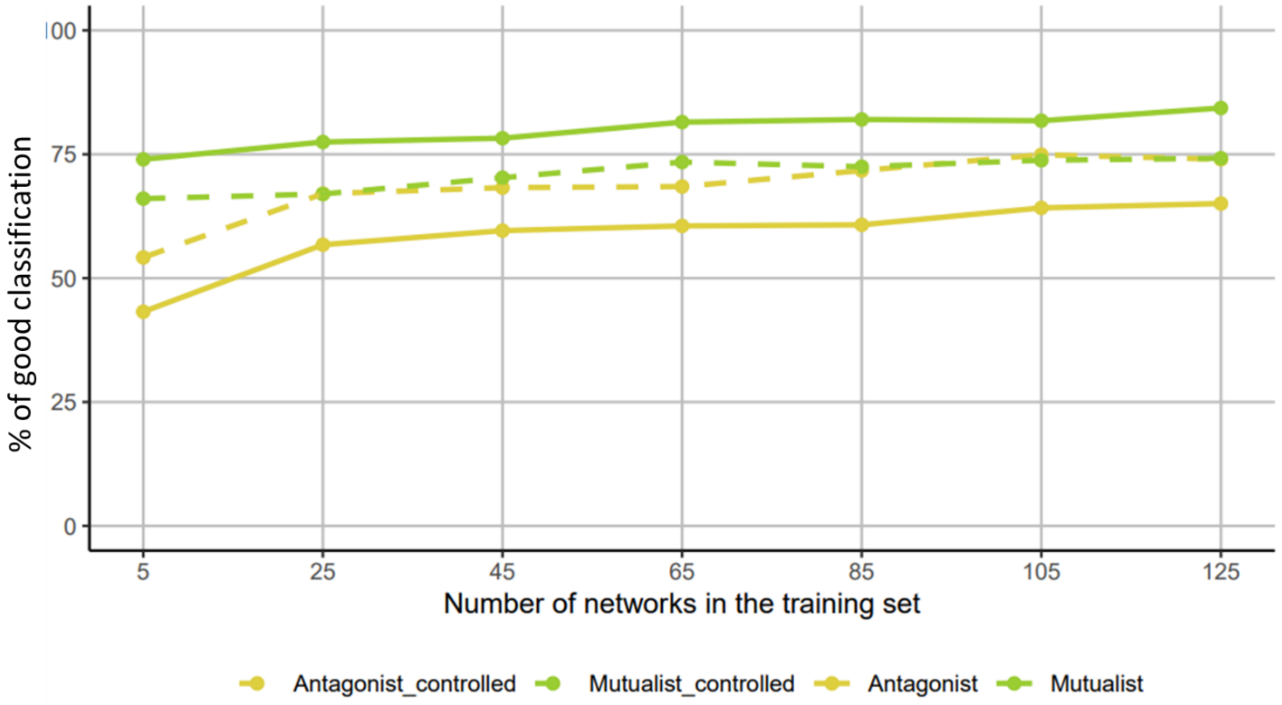

% of good classification
100
75
50
25
0
5
25
45
65
85
105
125
Number of networks in the training set
Antagonist_controlled
Mutualist_controlled
Antagonist
Mutualist


B

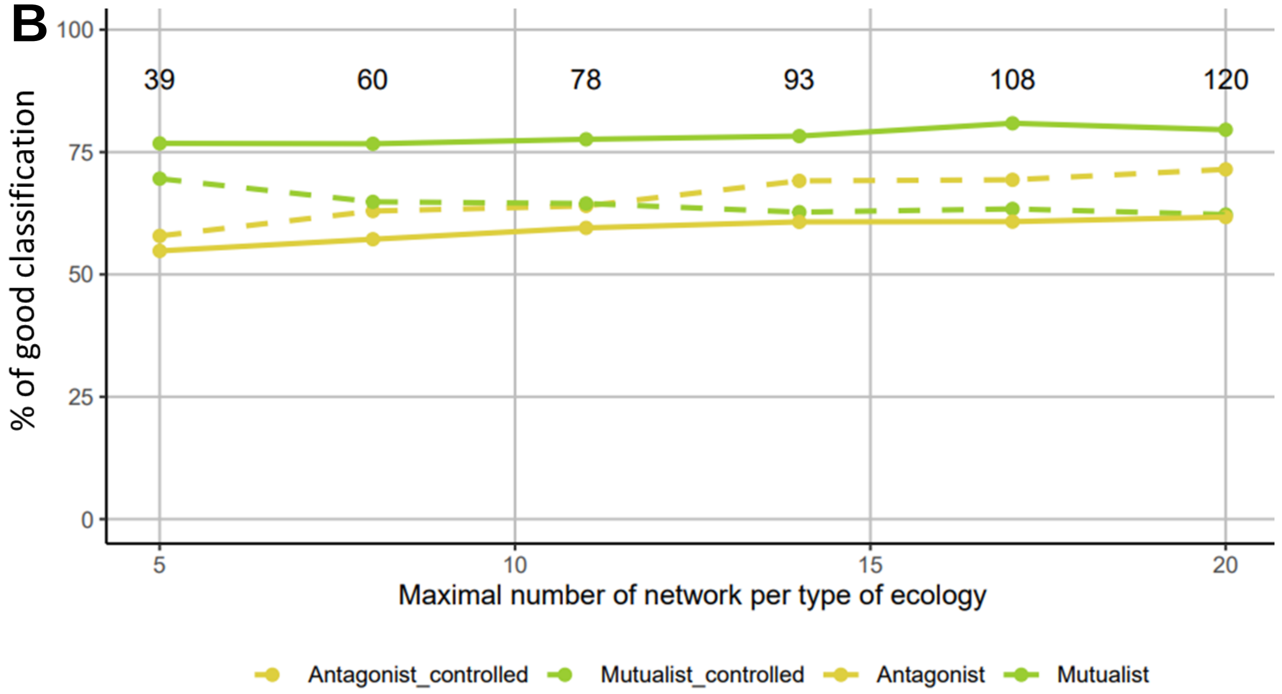

% of good classification
100
75
50
25
0
39
60
78
93
108
120
5
10
15
20
Maximal number of network per type of ecology
Antagonist_controlled
Mutualist_controlled
Antagonist
Mutualist

**Supplementary Figure 10: Global metrics of empirical and simulated networks.**

The panels represent the connectance (A), the nestedness (B; NODFc), the modularity (C; Newman's index), the difference in guilds size (D), and the sum (E) of guilds size for simulated (BipartiteEvol) and empirical networks. Asterisks indicate the statistical significance associated with a Wilcoxon-Mann-Whitney test performed to compare simulated and empirical networks on the different global metrics (*** = p-value<0.001, **=p-value<0.01).

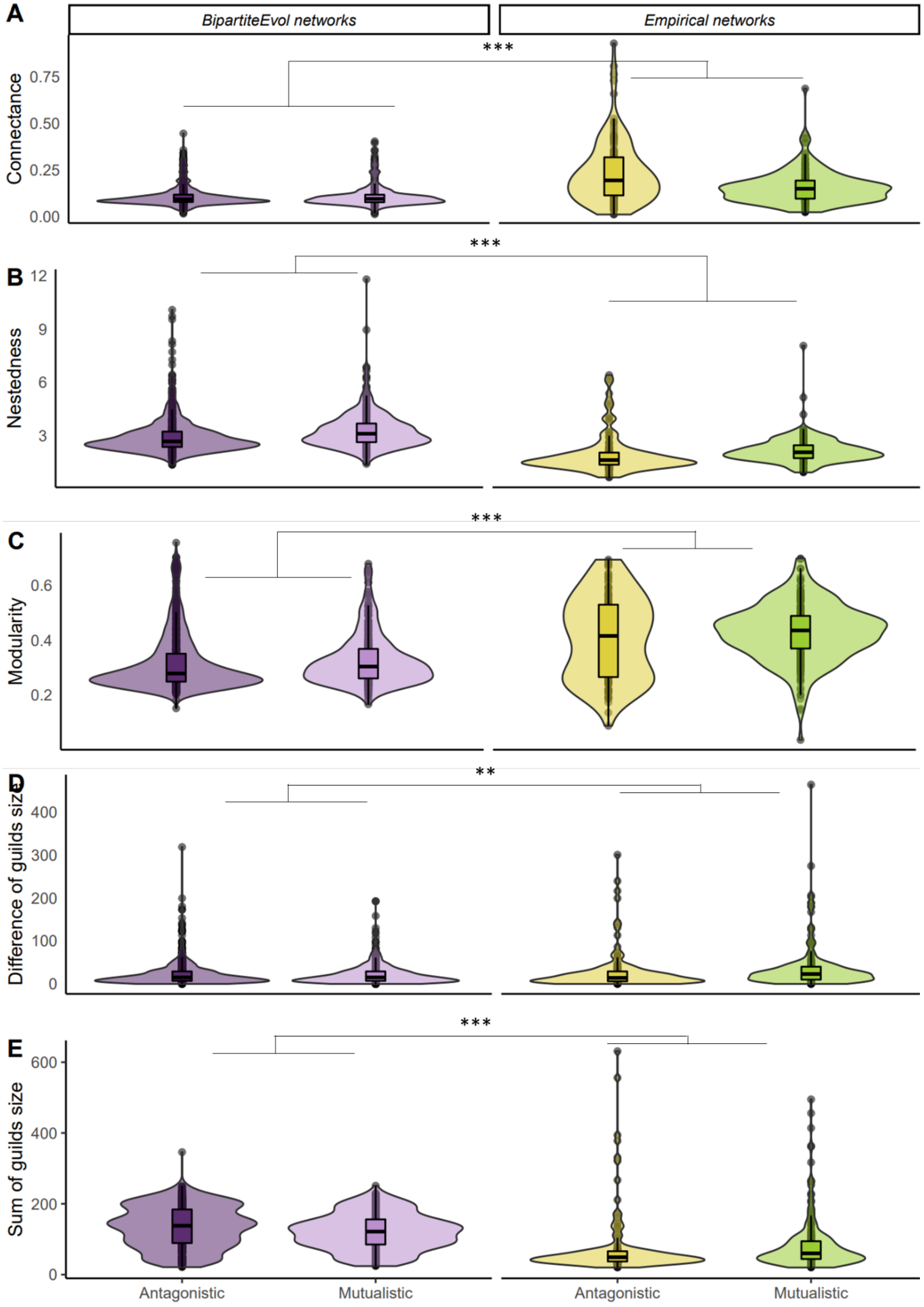
A
BipartiteEvol networks
Empirical networks
***
Connectance
0.75
0.50
0.25
0.00
B
***
Nestedness
12
9
6
3
C
***
Modularity
0.6
0.4
0.2
D
**
Difference of guilds size
400
300
200
100
0
E
***
Sum of guilds size
600
400
200
0
Antagonistic
Mutualistic
Antagonistic
Mutualistic

**Supplementary Figure 11: Principal coordinate analysis (PCA) performed on the global metrics of the empirical and simulated networks shows realistic BipartiteEvol simulations in terms of global metrics.**

PCA was performed using global metrics (nestedness, modularity, connectance, and the two metrics of network sizes) and we represented the projection on the two principal components (PC1 and PC2) for both empirical and simulated networks (A). For presentation purposes, we sampled 300 BipartiteEvol networks. The percentage of explained variance of the PCA is 40.9 % and 24.4 %.

Note that on PCA, the BipartiteEvol networks that are outside of the range of empirical networks correspond to large networks with more species than observed in empirical networks.

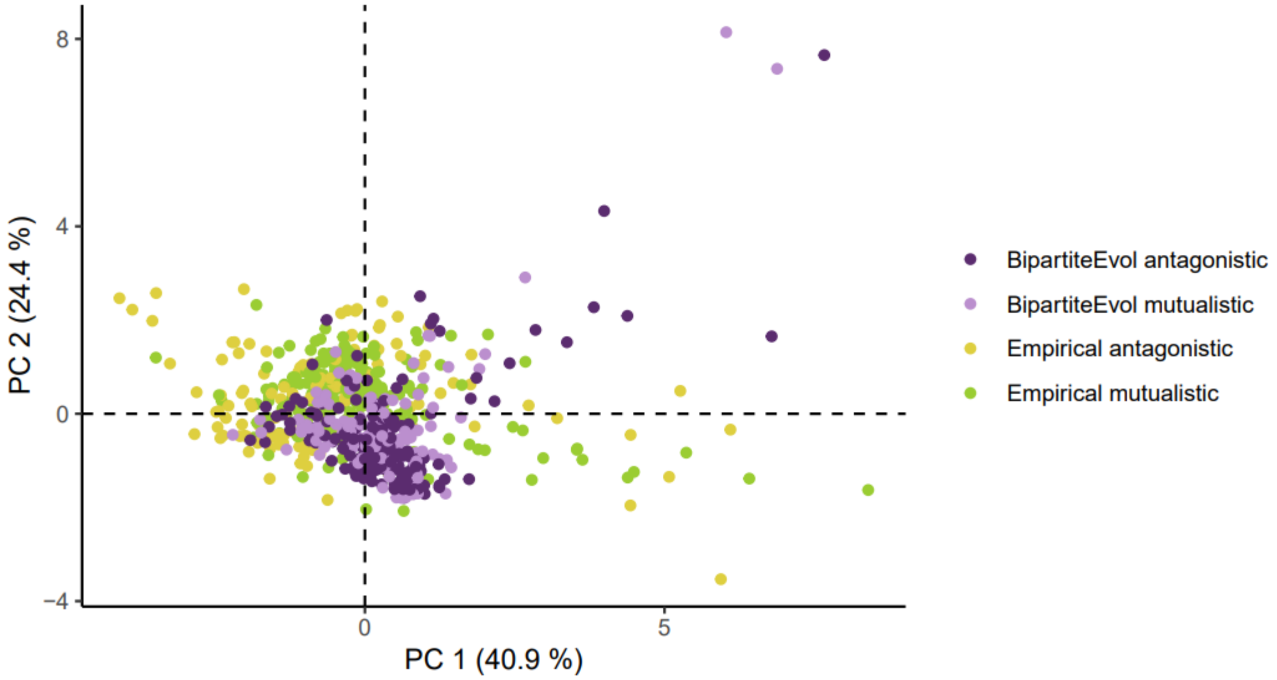

**Supplementary Figure 12: The principal coordinate analysis (PCA) performed on the Laplacian spectral density of both empirical and simulated networks shows that simulations failed at exploring the heterogeneity of empirical networks.**

PCA was performed using the Laplacian spectral density values and we represented the projection on the two principal components (PC1 and PC2) of both empirical and simulated networks. For presentation purposes, we sampled 300 BipartiteEvol networks. The percentage of explained variance is respectively 39.8 % and 11.6 % for the two first principal components of the PCA.

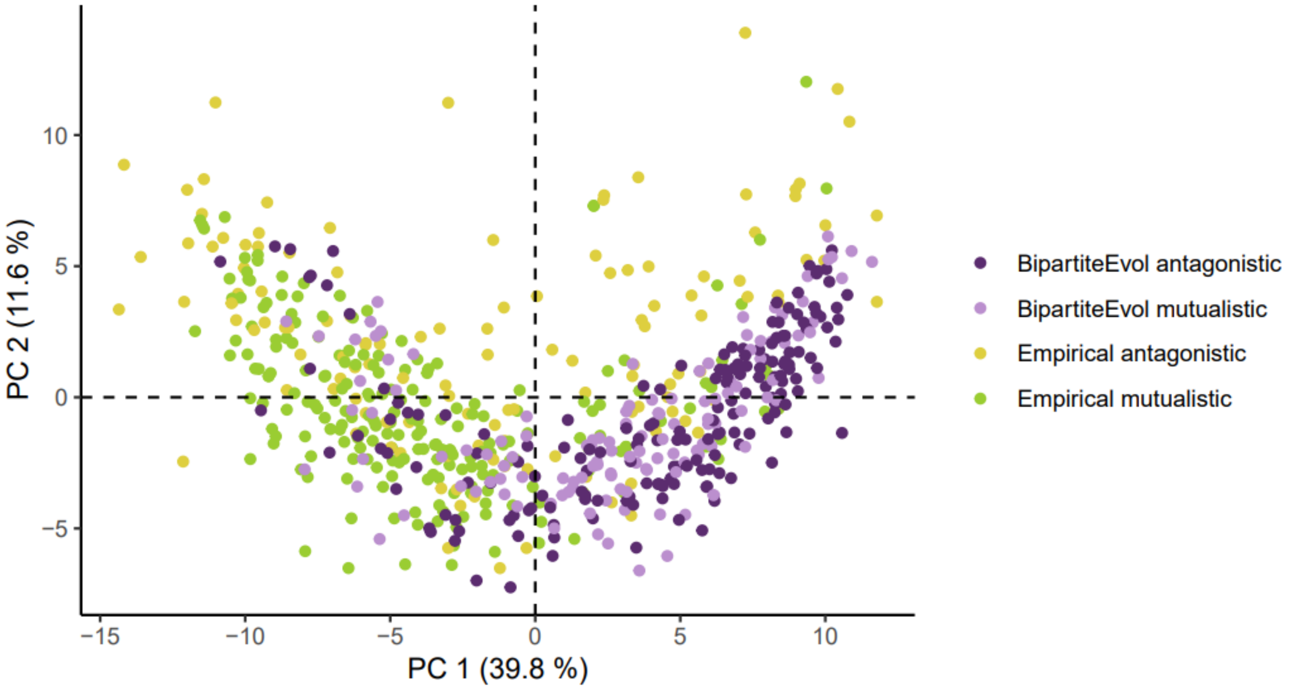

**Supplementary Figure 13: The principal coordinate analysis (PCA) performed on the motif frequencies of simulations and empirical networks shows that simulated networks failed at exploring the heterogeneity of empirical networks.**

PCA was performed using motif frequencies and we represented the projection on the two principal components (PC1 and PC2) of both empirical and simulated networks. For presentation purposes, we sampled 300 BipartiteEvol networks. The percentage of explained variance is respectively 25.9 % and 20.5% for the two first principal components of the PCA.

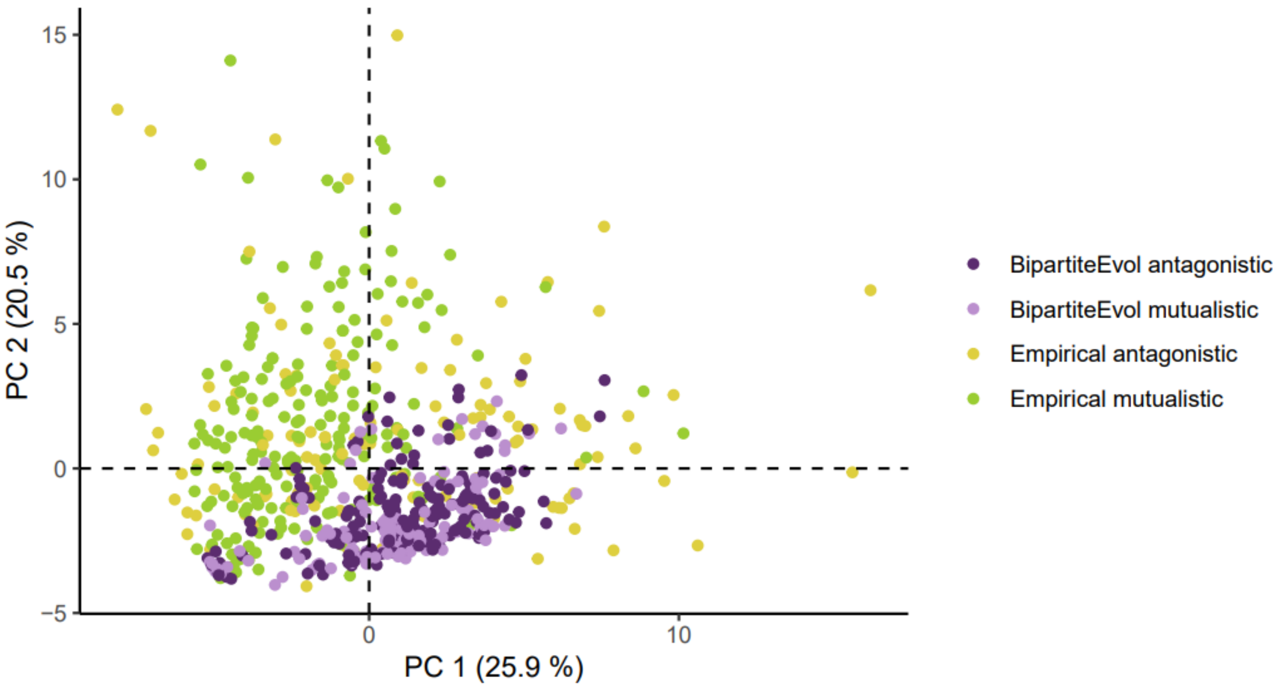

**Supplementary Figure 14: Some motif frequencies differ significantly between simulations and empirical networks.**

We compared the frequencies of all 44 motifs (A: 1 to 10; B: 11 to 20; C: 21 to 30; D: 31 to 44) between BipartiteEvol simulations (in purple), and empirical networks (yellow for mutualistic; green for antagonistic networks). Motifs starting from number 18 have 6 nodes, while others have fewer. Linear models were used for each motif with the type of network (simulated *versus* empirical) as the only effect (Fisher test: p-value<0.01). Boxplots present the median surrounded by the first and third quartiles, and whiskers extend to the extreme values but no further than 1.5 of the interquartile range.

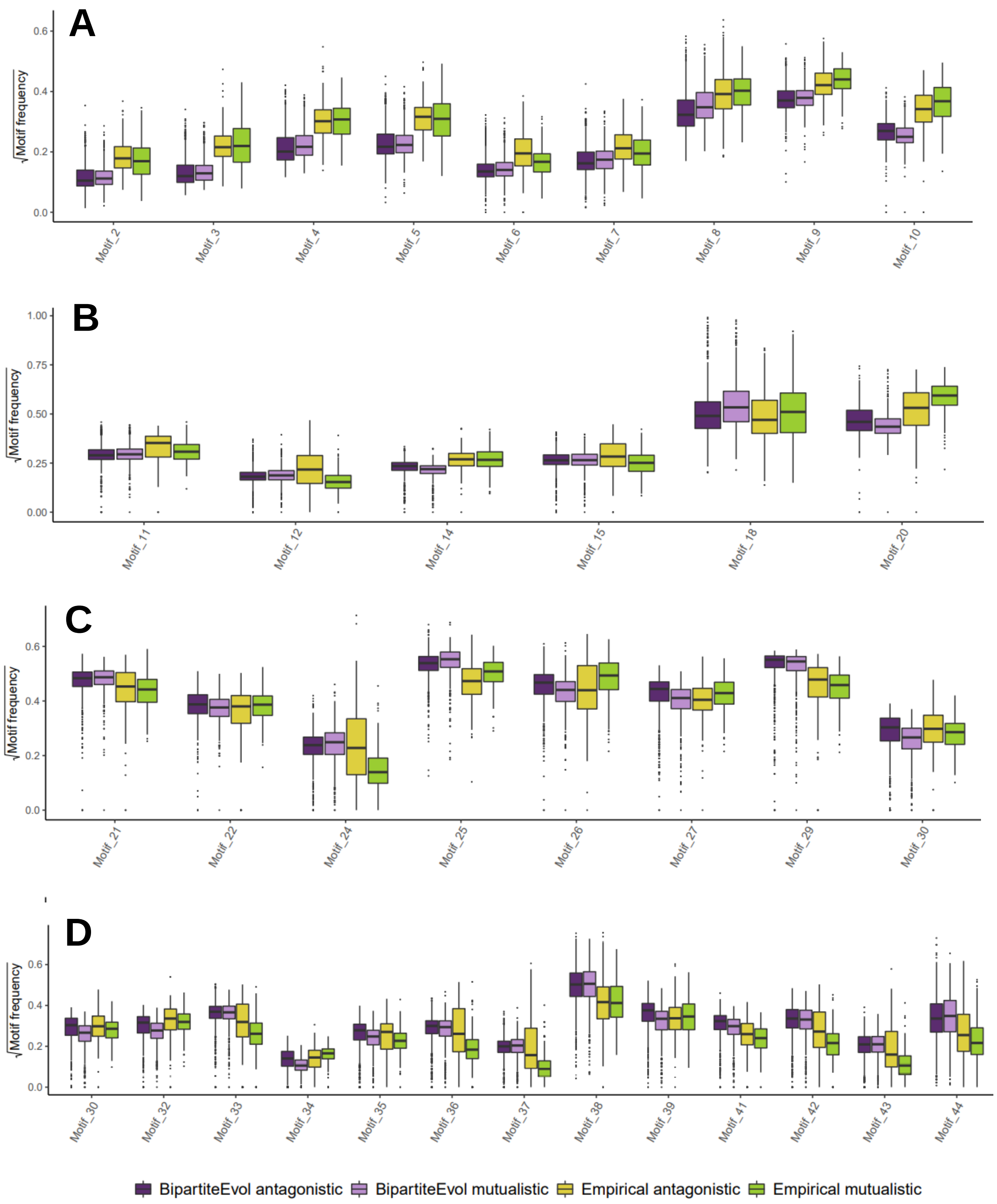
A
B
C
D
√Motif frequency
Motif_2
Motif_3
Motif_4
Motif_5
Motif_6
Motif_7
Motif_8
Motif_9
Motif_10
Motif_11
Motif_12
Motif_14
Motif_15
Motif_18
Motif_20
Motif_21
Motif_22
Motif_24
Motif_25
Motif_26
Motif_27
Motif_29
Motif_30
Motif_32
Motif_33
Motif_34
Motif_35
Motif_36
Motif_37
Motif_38
Motif_39
Motif_41
Motif_42
Motif_43
Motif_44
BipartiteEvol antagonistic
BipartiteEvol mutualistic
Empirical antagonistic
Empirical mutualistic

**Supplementary Figure 15: Differences between the Laplacian spectral densities of simulated BipartiteEvol and empirical networks.**

We compared the spectral density of the Laplacian matrix between BipartiteEvol simulations (in purple), and empirical networks (yellow for mutualistic; green for antagonistic networks) (A: spectral density of eigenvalues between 0 and 0.25; B: between 0.25 and 0.5; C: between 0.5 and 0.75; D: between 0.75 and 1). Linear models were used for each range of eigenvalues with the type of networks (simulated *versus* empirical) as the only effect (Fisher test: p-value<0.01). Boxplots present the median surrounded by the first and third quartiles, and whiskers extend to the extreme values but no further than 1.5 of the interquartile range.

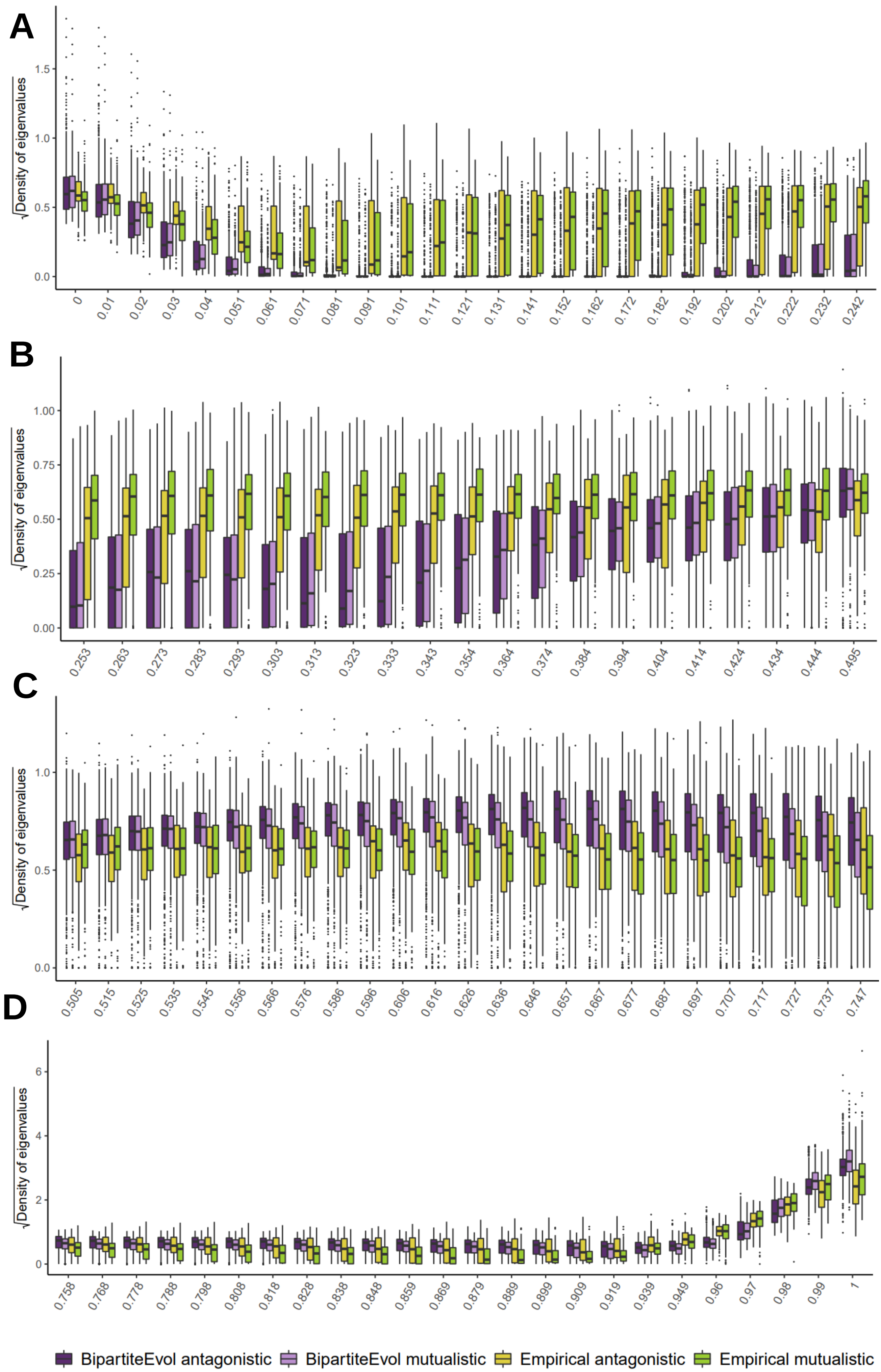

A
B
C
D
√Density of eigenvalues
0.0
0.5
1.0
1.5
0
0.01
0.02
0.03
0.04
0.051
0.061
0.071
0.081
0.091
0.101
0.111
0.121
0.131
0.141
0.152
0.162
0.172
0.182
0.192
0.202
0.212
0.222
0.232
0.242
0.00
0.25
0.50
0.75
1.00
0.253
0.263
0.273
0.283
0.293
0.303
0.313
0.323
0.333
0.343
0.354
0.364
0.374
0.384
0.394
0.404
0.414
0.424
0.434
0.444
0.495
0.505
0.515
0.525
0.535
0.545
0.556
0.566
0.576
0.586
0.596
0.606
0.616
0.626
0.636
0.646
0.657
0.667
0.677
0.687
0.697
0.707
0.717
0.727
0.737
0.747
0
2
4
6
0.758
0.768
0.778
0.788
0.798
0.808
0.818
0.828
0.838
0.848
0.859
0.869
0.879
0.889
0.899
0.909
0.919
0.939
0.949
0.96
0.97
0.98
0.99
1
BipartiteEvol antagonistic
BipartiteEvol mutualistic
Empirical antagonistic
Empirical mutualistic

**Supplementary Figure 16: Classification accuracies drop when randomizing the empirical networks, indicating that the good classification accuracy we obtained cannot be explained only by differences in network sizes, connectances, or degree distributions between antagonistic and mutualistic networks.**

We first trained the ANN with 80% of the empirical networks. Then, for the 20% remaining, we performed network randomization while keeping the network size fixed, and either the connectance fixed, or the degree distribution fixed. We tested the trained ANN on these randomized networks. We repeated this randomization strategy so that each empirical network was randomized 200 times and extracted (a) the mean percentage of correct classifications and (b) the mean F-score. Boxplots present the median of these 200 randomizations surrounded by the first and third quartiles, and whiskers extend to the extreme values. The horizontal lines correspond to (a) the percentage of correct classifications and (b) the F-scores of non-randomized networks (S4 Table). (c) We also indicate the projection of the motif frequencies of both empirical (yellow or green points) and randomized (grey and black; using a subset of all randomizations) networks in the two first components of a PCA. (d) Distribution of the distances of each empirical network to the PCA-centroid of its randomized networks with fixed connectance or degree distribution. We indicate the result of a Wilcoxon rank sum test between the two distributions (for randomizations with fixed connectance and size: W= 14884, p-value = 0.18; for randomizations with fixed degree and size: W=17220, p-value<0.001). (e) Overlap between the volumes occupied by empirical networks and their randomized counterparts using the Jaccard similarity index.

This analysis suggests that differences in network sizes, connectances, and degree distributions between antagonistic and mutualistic networks may help classify networks (percentages of correct classification and F-scores are not random at 50/50), but are not enough to explain the high classification accuracies we obtained.

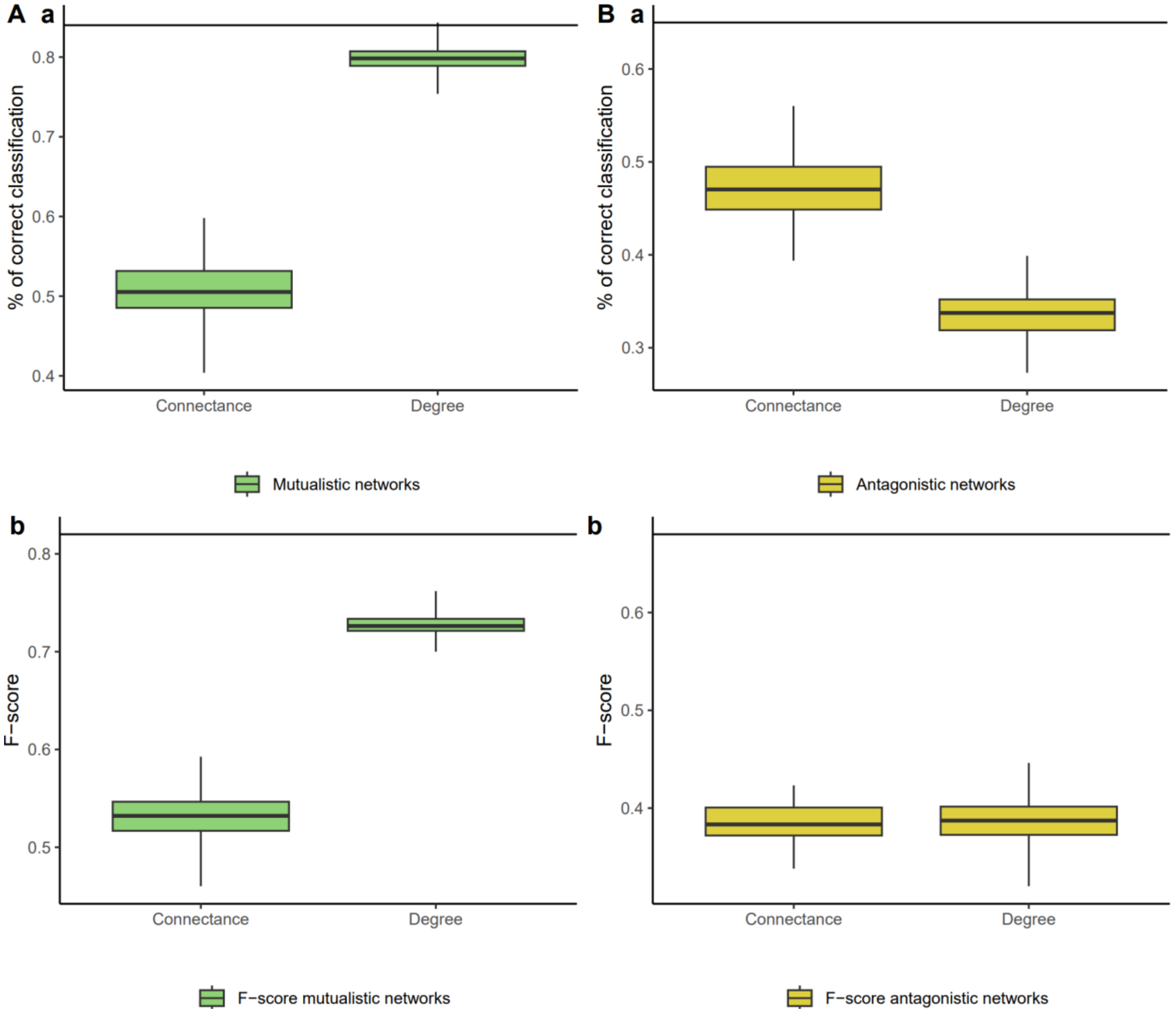
A a
% of correct classification
0.8
0.7
0.6
0.5
0.4
Connectance
Degree
Mutualistic networks
B a
% of correct classification
0.6
0.5
0.4
0.3
Connectance
Degree
Antagonistic networks
b
F−score
0.8
0.7
0.6
0.5
Connectance
Degree
F−score mutualistic networks
b
F−score
0.6
0.5
0.4
Connectance
Degree
F−score antagonistic networks

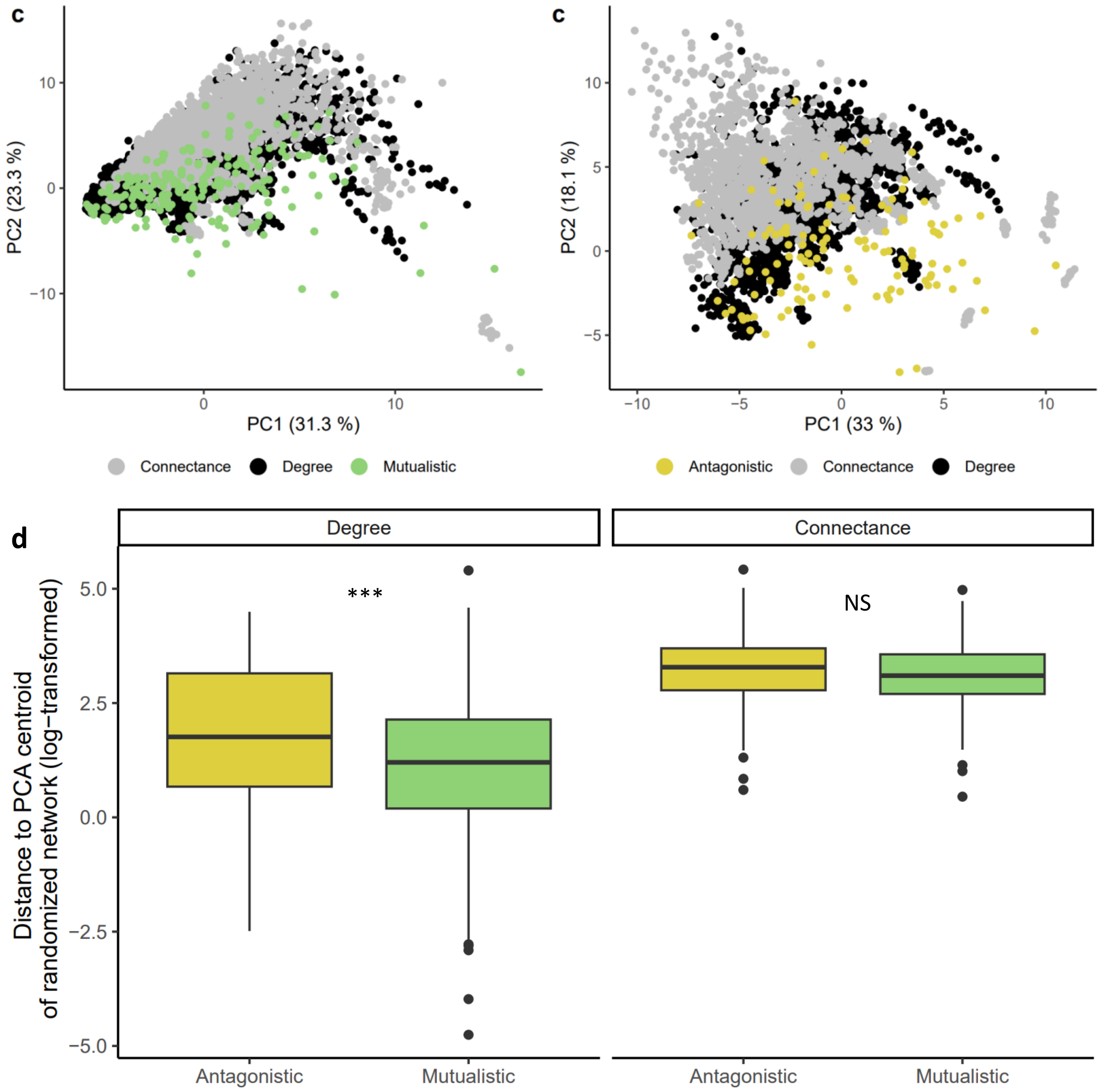


| e | Type of randomization | Antagonistic | Mutualistic |
|---|---|---|---|
| Overlapp in the PCA space between orginial and simulated networks (Jaccard similarity) | Degree | 0.419 | **0.472** |
| | Connectance | 0.34 | **0.397** |

**Supplementary Figure 17: The high classification accuracy we obtained is not driven only by the signal from the type of ecology.**

We performed randomization analyses were we randomly shuffled the interaction type (antagonistic versus mutualistic) of the different types of ecology (e.g. pollination networks were assigned as "antagonistic" and parasitic networks as "mutualistic") while maintaining 4 (*resp.* 5) different types of ecology for antagonistic (*resp.* mutualistic) networks. Using the motif frequencies of networks in this randomized dataset, we trained the ANN with 80% of the networks and tested it on the remaining 20%. We repeated this 50 times and recorded the mean percentage of correct classifications. We repeated this training-testing process for 200 shuffled datasets.

We present here (a) the mean percentage of correct classifications and F-scores as well as (b) the difference in percentage of correct classification and F-scores between antagonistic and mutualistic networks as a function of the number of networks which type of interaction was changed upon shuffling. Each point corresponds to a randomization. Note that when nearly all the networks are shuffled (high values on the x-axis), the type of ecologies of antagonistic (*resp.* mutualistic) networks are being replaced by the ones from mutualistic (*resp.* antagonistic) networks, which explains the symmetry of the curves. The interpretation of the signal from the type of ecology can be made when half of the networks are shuffled. We see that shuffling the type of interaction of each type of ecology decreases the average percentages of correct classifications (*resp.* F-scores) by about 10% (*resp.* 8%) while increasing the differences in percentage of correct classification (*resp.* F-scores) between antagonistic and mutualistic networks by about 30% (*resp.* 0.3). When shuffling the type of ecology the ANN cannot distinguish antagonistic from mutualistic networks anymore and it instead classifies most networks as one of the two interaction types.

In rare occasions, we noticed that some randomizations performed better than what is reported in Table S4. This can for instance happen when herbivory networks are attributed the mutualistic type: in the original classification (i.e. without shuffling), herbivory networks tend to be more frequently (wrongly) classified as mutualistic (see Table 1); hence, if they are attributed the antagonistic interaction type upon shuffling, both the percentage of correct classification and the F-scores are higher. In general however, shuffling interaction types drastically reduces both F-scores and percentage of correct classification, demonstrating that there is a signal of interaction type beyond ecology.

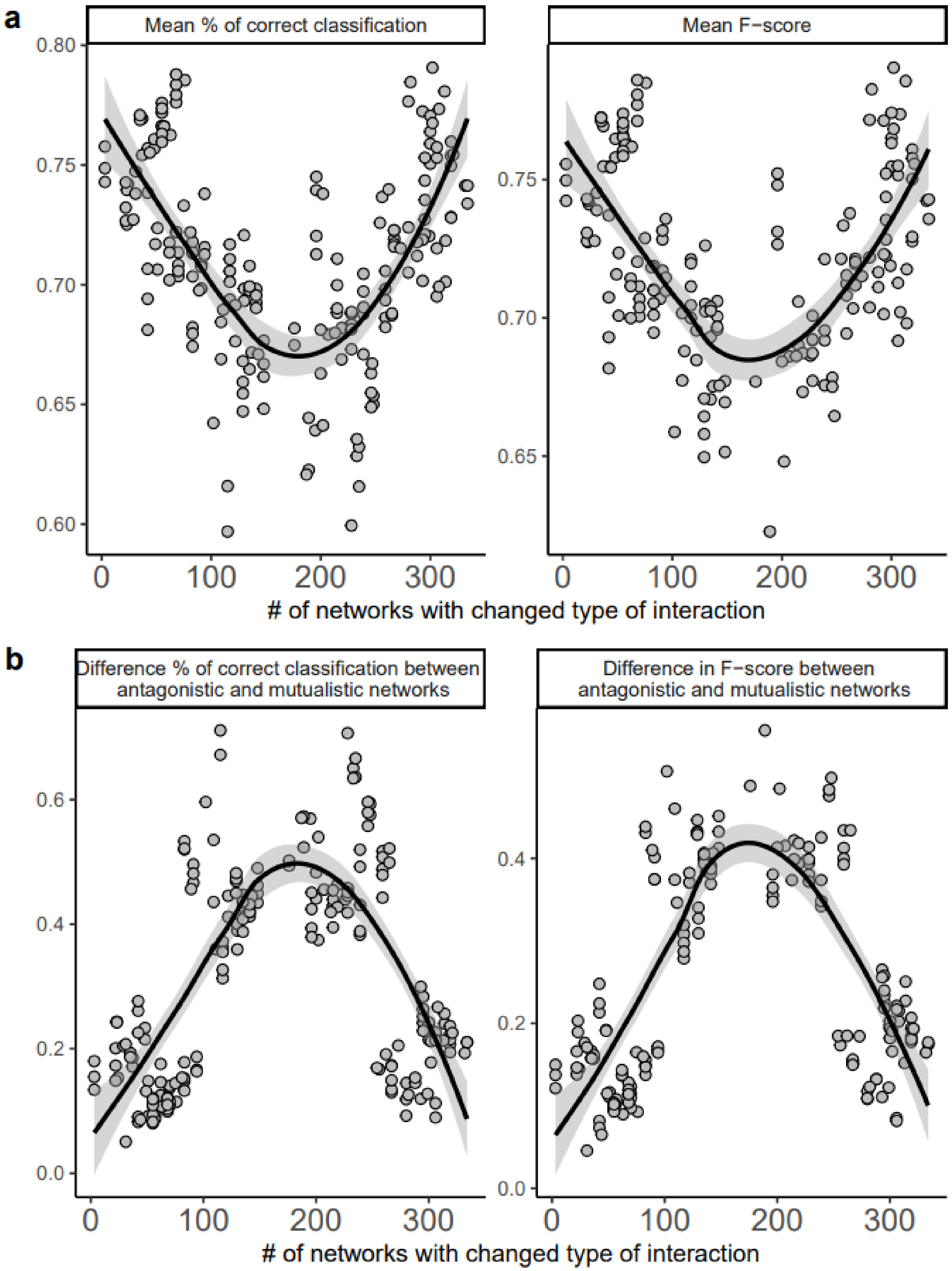

**Supplementary Figure 18: Distributions of the percentages of correct classification of each network in the dataset over 10,000 different training-testing sets.**

We performed 10000 different training-testing processes and computed the percentage of correct classification of each mutualistic (left panel) and antagonistic (right panel) network. We also indicate the boundaries between 'low' and 'high' classification confidence defined for Table 1 with vertical lines. (a) Colored by the type of interaction. (b) Colored by the type of ecology.

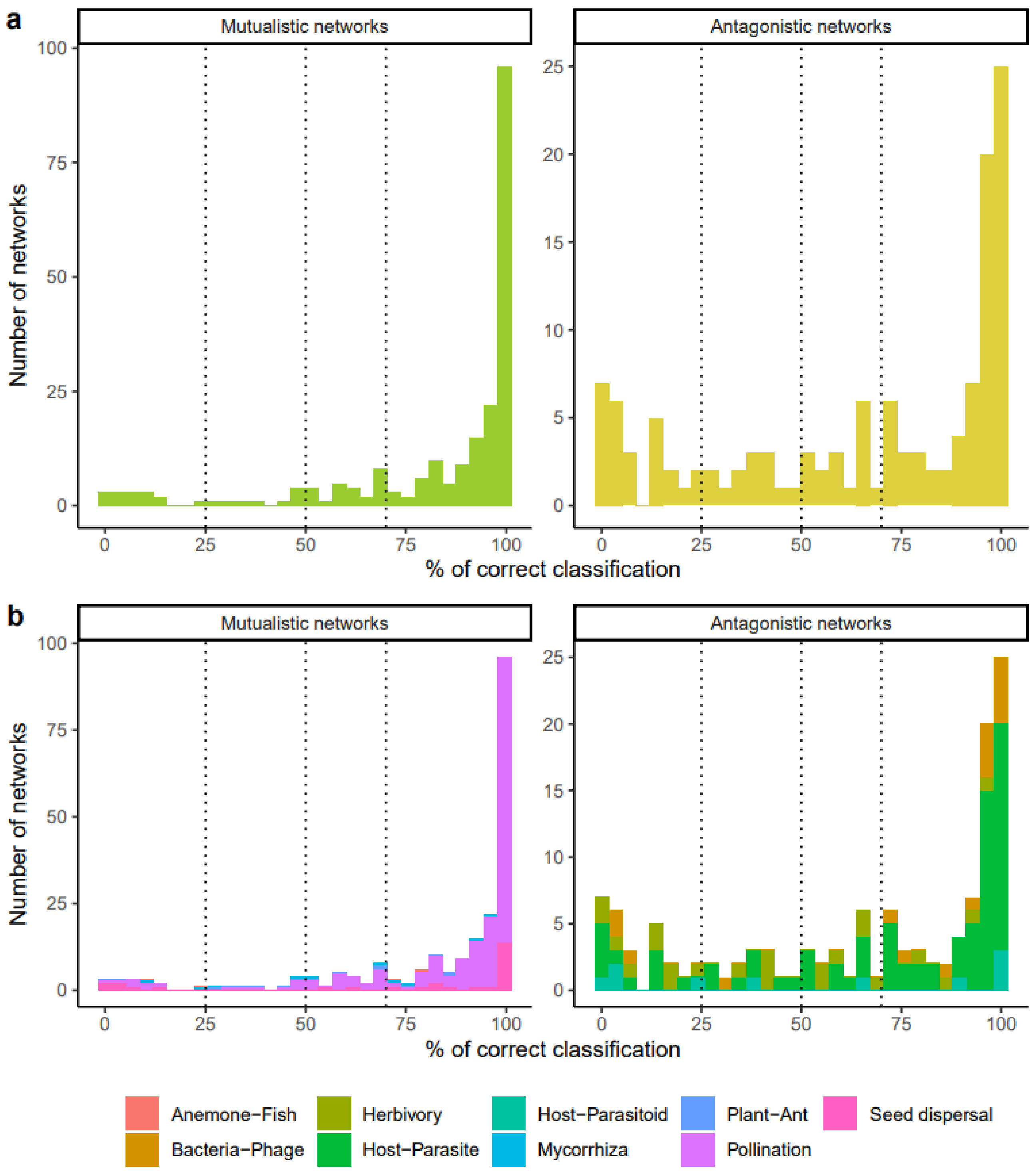
